# The Impact of Oxytocin on Neurite Outgrowth and Synaptic Proteins in *Magel2*-Deficient Mice

**DOI:** 10.1101/2020.09.23.309955

**Authors:** Alexandra Reichova, Fabienne Schaller, Stanislava Bukatova, Zuzana Bacova, Françoise Muscatelli, Jan Bakos

**Author notes:** Corresponding author: Jan Bakos, PhD., Institute of Experimental Endocrinology, Biomedical Research Center, Slovak Academy of Sciences, Dubravska cesta 9, 845 05, Bratislava, Slovakia.

## Abstract

Oxytocin contributes to the regulation of cytoskeletal and synaptic proteins and could therefore affect the mechanisms of neurodevelopmental disorders, including autism. Both the Prader-Willi syndrome and Schaaf-Yang syndrome exhibit autistic symptoms involving the *MAGEL2* gene. *Magel2*-deficient mice show a deficit in social behavior that is rescued following postnatal administration of oxytocin. Here, in *Magel2*-deficient mice, we showed that the neurite outgrowth of primary cultures of immature hippocampal neurons is reduced. Treatment with oxytocin, but not retinoic acid, reversed this abnormality. In the hippocampus of *Magel2*-deficient pups, we further demonstrated that several transcripts of neurite outgrowth-associated proteins, synaptic vesicle proteins, and cell-adhesion molecules are decreased. In the juvenile stage, when neurons are mature, normalization or even overexpression of most of these markers was observed, suggesting a delay in the neuronal maturation of *Magel2*-deficient pups. Moreover, we found reduced transcripts of the excitatory postsynaptic marker, *Psd95* in the hippocampus and we observed a decrease of PSD95/VGLUT2 colocalization in the hippocampal CA1 and CA3 regions in *Magel2*-deficient mice, indicating a defect in glutamatergic synapses. Postnatal administration of oxytocin upregulated postsynaptic transcripts in pups; however, it did not restore the level of markers of glutamatergic synapses in *Magel2*-deficient mice. Overall, *Magel2* deficiency leads to abnormal neurite outgrowth and reduced glutamatergic synapses during development, suggesting abnormal neuronal maturation. Oxytocin stimulates the expression of numerous genes involved in neurite outgrowth and synapse formation in early development stages. Postnatal oxytocin administration has a strong effect in development that should be considered for certain neuropsychiatric conditions in infancy.

## Introduction

Recent studies have shown that multiple components of the neuronal cytoskeleton, including microtubule-associated proteins, cell-adhesion molecules, and small GTPases participate in the regulation of neurite outgrowth and synapse formation (Zeidán-Chuliá et al., 2013; Hart & Hobert, 2018; Cao, Deng & Pocock 2019; Ruhl et al., 2019). Abnormalities in neuritogenesis, elongation of axons and dendrites along with alterations in the expression of synapse-associated and microtubule-associated proteins might contribute to the structural basis of autism pathology (Bakos et al., 2015; Zatkova et al., 2016; Gilbert & Man, 2017).

Oxytocin (OT) plays a pleiotropic role from birth at the cellular level via binding to G-protein coupled receptors and triggering different signaling pathways (Busnelli & Chini, 2018; Jurek & Neumann, 2018). Oxytocin receptors (OXTR) are expressed in the hippocampal excitatory pyramidal neurons and inhibitory interneurons; both are known to be involved in the regulation of social behavior (Lin et al., 2017; Raam et al., 2017; Meira et al., 2018; Young & Song, 2020). Moreover, it has been suggested that OT controls the development of hippocampal excitatory neurons (Ripamonti et al., 2017). We have recently shown that neuritogenesis is under the control of OXTR (Lestanova et al., 2016; Zatkova et al., 2018). Furthermore, activation of the OXTRs modulates the expression of synaptic adhesion molecules in neuronal cell cultures (Zatkova et al., 2019). It is likely that OT could be involved in the processes of neurite outgrowth and related signaling cascades, however, the downstream effectors are poorly understood. Alterations in OT and its receptors have been shown to cause autism spectrum disorder (ASD)-like phenotypes in rodent models with deficiencies in social function (Sebat et al., 2007; Muscatelli et al., 2018), and thus any alteration in neurite outgrowth may participate in these pathophysiological mechanisms.

The Prader–Willi Syndrome (PWS) and the closely-related Schaaf-Yang Syndrome (SYS) are complex genetic syndromes, which manifest various autistic symptoms. ASD symptoms are more frequent in SYS patients (approx. 80%) who present point mutations in *MAGEL2* only, as compared to PWS (approx. 30 %) patients, who have a large chromosomal deletion including *MAGEL2* (Schaaf et al., 2013; Urreizti et al., 2017). To date, two mouse models deficient for *Magel2* have been established (Kozlov et al., 2007; Schaller et al., 2010) and show an impairment of social behavior and cognition at adulthood (Meziane et al., 2015; Fountain et al., 2017). In one of these models, the neonatal treatment of *Magel2^tm1.1Mus^* mice with OT has been sufficient for the prevention of social and learning deficits in adulthood, in particular social memory (Schaller et al., 2010; Meziane et al., 2015, Bertoni et al., 2020). More specifically, in *Magel2*-deficient mice, we revealed cellular, physiological and biochemical alterations of hippocampal regions, which are known to be associated with social memory engrams involving the OT-system; those alterations are corrected following perinatal administration of OT (Bertoni et al., 2020).

In the present study, we hypothesize that 1) neurite outgrowth is altered in primary cultures of immature hippocampal neurons isolated from *Magel2^tm1.1Mus^* (hereafter named *Magel2^-/-^*) deficient pups, 2) *Magel2* deficiency impairs the regulation of hippocampal neurite outgrowth-associated and synaptic transcripts in early postnatal development and 3) OT treatment could restore normal neurite outgrowth and synaptic formation in the early development of *Magel2*-deficient mice. In addition to the neuronal primary cultures, we also investigated two postnatal critical developmental stages: infancy (P5) and juvenile (P30), which are associated with non-matured (P5) or matured (P30) hippocampal neurons.

## Material and Methods

### Animals

Mice were handled and cared for in accordance with the Guide for the Care and Use of Laboratory Animals (N.R.C., 1996) and the European Communities Council Directive of September 22nd, 2010 (2010/63/EU, 74). Experimental protocols were approved by the institutional Ethical Committee guidelines for animal research with the accreditation no. B13-055-19 from the French Ministry of Agriculture. We maintained grouped-house mice (3-5 mice/cage). All efforts were made to minimize the number of animals used. To overcome the heterogeneity in the analysis of the heterozygous^+m/-p^ *Magel2^tm1.1Mus^* mice (due to the stochastic expression of the maternal allele of *Magel2* when the paternal allele is deleted (Matarazzo & Muscatelli, 2013), we used here *Magel2^tm1.1Mus^* homozygous^-/-^ mice, (hereafter named *Magel2^-/-^* deficient mice) generated as previously described (Schaller et al., 2010). Mice colonies were maintained in the C57BL/6J background. Due to the parental imprinting of *Magel2* (paternally expressed only), in order to obtain heterozygote mice (+m/-p), males carrying the mutation on the maternal allele (-m/+p) were crossed with wild-type C57BL/6J females. To obtain homozygote mice (-m/-p), *Magel2*-deficient homozygote males and females were crossed. It is also important to note that *Magel2^-/-^* mothers were observed as having similar maternal behavior as the wild-type (WT). All mice were genotyped by PCR stemming from DNA extracted from tail snips (around 3 mm) and using the primers shown in Table 1.

**Table 1:**
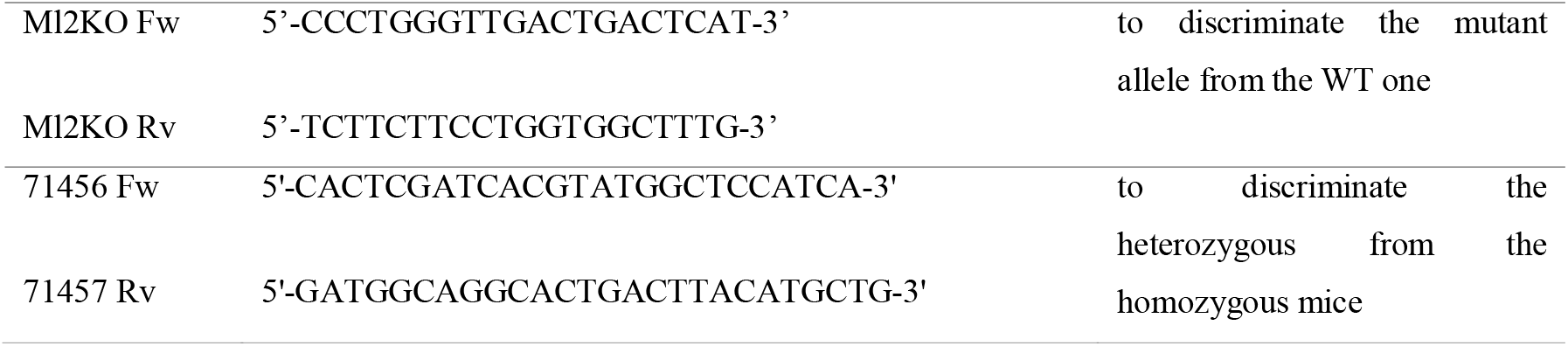
Primer sequences for genotyping. Fw – forward; Rv – reverse.

### Preparation of Primary Hippocampal Neurons

Brains from WT (n=15 pups/morphological analysis, n=9 pups/quantification of excitatory synaptic proteins) or *Magel2^-/-^* mice (n=7 pups/morphological analysis, n=9 pups/ quantification of excitatory synaptic proteins) at postnatal day 0 (P0) were dissected on icecold Hank’s Balanced Salt Solution – HBSS (Gibco, USA), supplemented with 1% Antibiotic-Antimycotic (Gibco, USA) and 0.3 M HEPES (Gibco, USA) under a stereomicroscope in order to collect hippocampi. The hippocampal tissues were dissociated for 20 min at 37°C with an enzymatic solution (HBSS, 0.1% Trypsin, 0.1mg/ml DNAse I). Cells were resuspended and plated on 24-well plates containing cover slips coated with 10 μg/ml poly-D-lysine (Sigma-Aldrich, Germany) at a density of 0.8×10^5^/ml in RPMI medium (Sigma-Aldrich, Germany) containing 10% fetal bovine serum. After 3h of cell plating in a 37°C and 5% CO_2_ incubator, the medium was replaced with a selective growing medium: Neurobasal A (Gibco, USA), 1% Antibiotic-Antimycotic (Gibco, USA), 1% Glutamax (Gibco, USA) and 2% supplement B27 (Invitrogen, USA); day *in vitro* 1 (DIV1). After 5 days *in vitro*, 50% of growing media were exchanged. For the purpose of the evaluation of neurite outgrowth and quantification of excitatory synaptic proteins in immature neurons, 1 μM OT (Bachem, Switzerland) or 10 μM all-trans retinoic acid -ATRA (Sigma-Aldrich, Germany) were added to the medium at DIV7 and DIV8 (48h treatment).

### Immunocytochemistry

After 48 h incubation (DIV9), the medium was removed and primary hippocampal cells were fixed with 4% AntigenFix (Diapath, Italy), pH 7.4 for 20 min at room temperature (RT). The coverslips were washed 2 times with cold PBS and blocked in 3% (v/v) normal goat serum (NGS) / donkey serum (DS) and 0.1% Triton X-100 for 30 min at RT. Cells were stained with primary antibodies diluted in PBS with 3% serum, 0.1% Triton X-100 (Table 2) during 2h at RT. Afterwards, the coverslips were rinsed 3 times with cold PBS and incubated with corresponding fluorescent secondary antibodies diluted in PBS (Table 3) for 1 h at RT. The nuclei were stained with 300 nM DAPI (4’,6-diamidino-2-phenylindole; Thermo Fisher Scientific, Slovakia) for 1 min. Coverslips were mounted on glass slides with Fluoromount-G (Sigma-Aldrich, Germany).

**Table 2:** Primary antibodies. ICC – Immunocytochemistry; IHC – Immunohistochemistry; WB – Western Blot Analysis; Ras homolog family member - RHO(A,B); Postsynaptic density protein 95 - PSD95; Glyceraldehyde 3-phosphate dehydrogenase – GAPDH; Microtubule Associated Protein 2 - MAP2; Vesicular Glutamate Transporter 2 – VGLUT2; Glutamic Acid Decarboxylase – GAD(65/67).

| *Name* | *Host species* | *Dilution* | *Method* | *Product number* |
| --- | --- | --- | --- | --- |
| **anti-RHOA** | Rabbit | 1:1000 | WB | # 2117S; Cell Signaling, USA |
| **anti-RHOB** | Rabbit | 1:500 | WB | AB_155149; Abcam, UK |
| **anti-PSD95** | Goat | 1:500 | WB/IHC/ICC | AB_12093; Abcam, UK |
| **anti-GAPDH** | Mouse | 1:1000 | WB | G8795; Sigma-Aldrich, Germany |
| **anti-GAPDH** | Rabbit | 1:1000 | WB | G9545; Sigma-Aldrich, Germany |
| **anti-VGLUT2** | Chicken | 1:5000 | IHC/ICC | 135 416; Synaptic Systems, Germany |
| **anti-MAP2** | Mouse | 1:2000 | ICC | M4403; Sigma-Aldrich, Germany |
| **anti-GAD65+GAD67** | Rabbit | 1:1000 | WB | ab11070; Abcam, UK |

**Table 3:** Fluorescent secondary antibodies. ICC – Immunocytochemistry; IHC – Immunohistochemistry; WB – Western Blot Analysis.

| *Name* | *Host species* | *Dilution* | *Method* | *Product number* |
| --- | --- | --- | --- | --- |
| **anti-rabbit**<br>CF<sup>TM</sup> 770 | Goat | 1:5000 | WB | SAB4600215; Sigma-Aldrich, Germany |
| **anti-mouse**<br>CF<sup>TM</sup> 680R | Donkey | 1:5000 | WB | SAB4600207; Sigma-Aldrich, Germany |
| **anti-mouse**<br>Alexa Fluor 488 | Goat | 1:500 | IHC/ICC | A-11001; Thermo Fisher Scientific, Slovakia |
| **anti-goat**<br>Alexa Fluor 680 | Donkey | 1:10 000 | WB | AB_175776; Abcam, UK |
| **anti-goat**<br>Alexa Fluor 555 | Donkey | 1:500 | IHC/ICC | A-21432; Thermo Fisher Scientific, Slovakia |
| **anti-chicken**<br>Alexa Fluor 647 | Goat | 1:500 | IHC/ICC | A-21449; Thermo Fisher Scientific, Slovakia |

#### A) Morphological Analysis

MAP2 (Microtubule Associated Protein 2) positive cells were considered neurons. Image capturing was performed using an Apotome2 microscope (Zeiss, Germany) at high magnification (20x objective, numerical aperture 0.8). Six coverslips per experimental group and at least 7 areas of interest per coverslip were evaluated by Fiji/ImageJ software. Arborization of dendritic trees was assessed by Sholl analysis plugin (Cloarec et al., 2019). Calculation of the number of intersecting dendrites with concentric circles in the interval from the soma up to 200 μm was performed. The length of the longest neurite was quantified from the edge of the nucleus to the apical end of the neurite by three independent members of the research team by manual tracing.

#### B) Quantification of excitatory synaptic proteins

For the analysis of excitatory synaptic proteins in primary hippocampal neurons, triple staining for the Postsynaptic Density Protein 95 (PSD95), Vesicular Glutamate Transporter 2 (VGLUT2) and MAP2 were performed. For image acquisition, a Nikon confocal microscope (Nikon ECLIPSE Ti-E, A1R+; Netherlands) was used at high magnification (40x objective, numerical aperture 1.3; resolution 1024 × 1024 pixels), and image analysis was carried out by Fiji/ImageJ software. Each experimental group consisted of 5 coverslips where at least 10 cells per coverslip were analyzed. Three individual dendritic segments (10 μm each) per neuron, situated at least 10 μm from the nucleus, were randomly chosen by two different researchers. Total fluorescence of PSD95 and VGLUT2 normalized on the MAP2 total fluorescence for the purpose of quantifying the excitatory synaptic proteins was evaluated.

### OT Treatment

Pregnant WT and *Magel2^-/-^* mice were monitored, and the day of birth was marked as postnatal day 0 (P0). The animals were sexed by anogenital distance and assigned to two neonatal treatment groups. Only the male mice were studied. Either a saline solution (20 μl) or 20 μl OT (Bachem, Switzerland) diluted in a saline (0.1 μg/μl) were given subcutaneously (s.c.) to the mice on days P2 and P3. The experimental protocol, which was previously used and described by Bakos et al. (2014), as well as a dose of OT was used according to the protocol of Schaller et al. (2010). On day P5, the mice were either 1) sacrificed by decapitation and the brains were quickly removed from the skull with the right/left hippocampi being separated, then deep frozen by liquid nitrogen and stored at −80°C or 2) deeply anesthetized by dormitor/zoletil and transcardially perfused with 4% AntigenFix (Diapath, Italy), pH 7.4. All brains were removed from the skull and postfixed in the same fixative overnight. After fixation, brain tissue was preserved in PBS with 0.01% sodium azide.

### Quantitative Real-Time PCR

Total RNA from the right hippocampi (n=6/group) was isolated by a phenol–chloroform method using TRI-reagent (MRC, Germany) according to the manufacturer’s protocol. The tissues separated from the P5 and P30 mice (WT and *Magel2^-/-^*) were analyzed. The concentration and purity of RNA was determined by a Nanodrop spectrophotometer (Thermo Fisher Scientific, Slovakia). The reverse transcription was carried out using a High-Capacity cDNA Reverse Transcription Kit (Thermo Fisher Scientific, Slovakia) according to the manufacturer’s instructions. Quantitative real-time PCR (qRT-PCR) was performed using Power SYBR^®^ Green PCR Master Mix (Thermo Fisher Scientific, Slovakia) and a QuantStudio 5 thermocycler (Thermo Fisher Scientific, Slovakia). Relative gene expression levels were calculated by using the Livak method (Livak & Schmittgen, 2001). The 2^-ΔΔCt^ value of each sample was calculated using *glyceraldehyde 3-phosphate dehydrogenase (Gapdh*) as a reference control gene (primers sequences in Table 4).

**Table 4:**
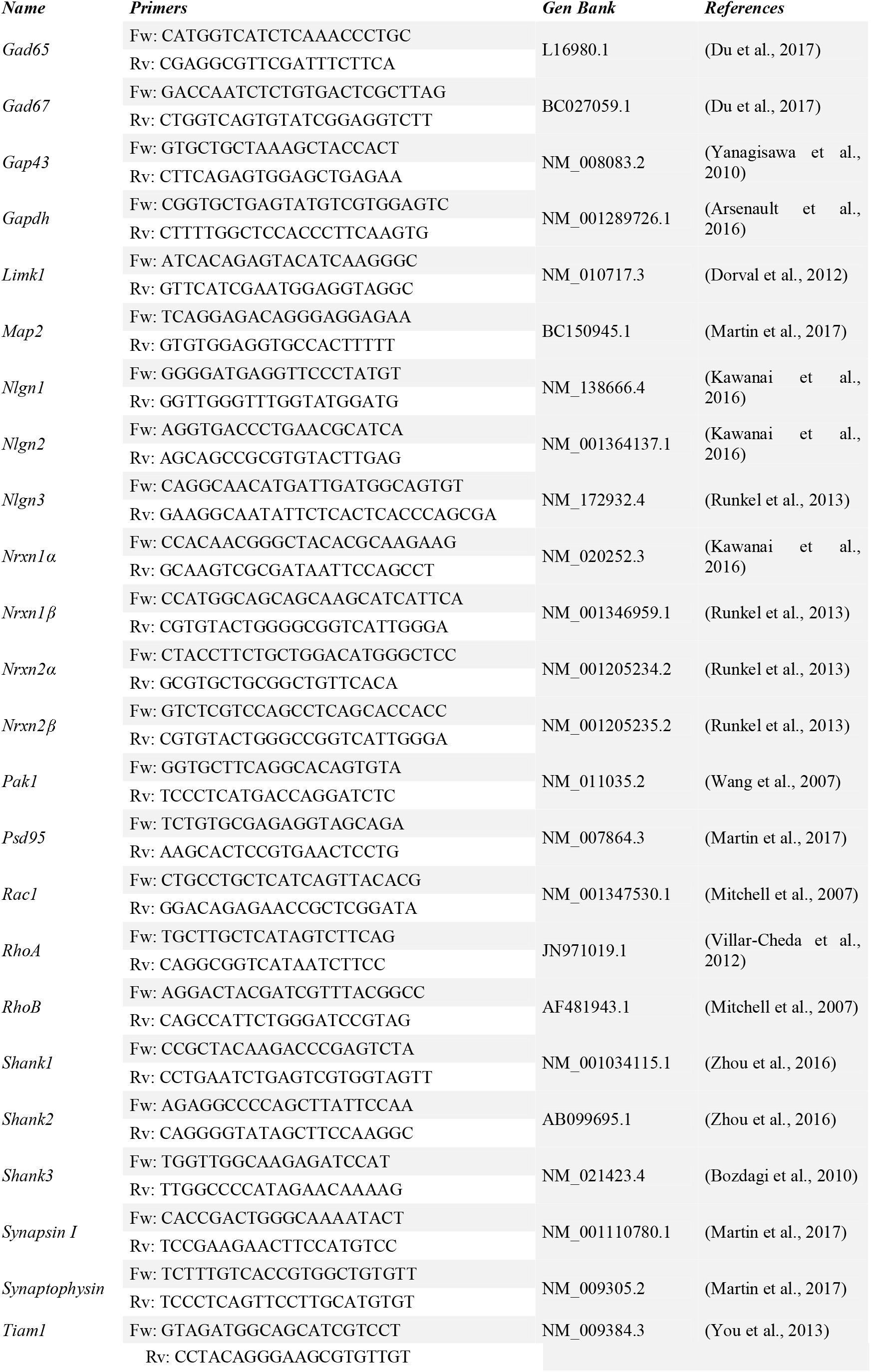
Primer sequences for qRT-PCR. Fw – forward; Rv – reverse; *Glyceraldehyde 3-phosphate dehydrogenase* – *Gapdh; Microtubule associated protein 2* – *Map2; Growth associated protein 43* - *Gap43; p21 activated kinase 1* - *Pak1; SH3 and multiple ankyrin repeat domains* - *Shank(1-3); Neuroligin* – *Nlgn(1-3); Neurexin* – *Nrxn(1α/β-2 α/β); Ras-related C3 botulinum toxin substrate 1* - *Rac1; T-cell lymphoma invasion and metastasis 1* - *Tiam1; LIM domain kinase 1* - *Limk1; Ras homolog family member* - *Rho(A,B); Glutamic acid decarboxylase* – *Gad(65,67); Postsynaptic density protein 95* - *Psd95*.

### Western Blot Analysis

The left hippocampi (n=6/group) from P5 and P30 mice (WT and *Magel2^-/-^*) were homogenized by gently shaking a tissue in a lysis buffer (100 mM NaCl, 10 mM Tris, 1 mM EDTA, 1 mM EGTA, 1% Triton X-100, 10% v/v glycerol, 0.1% sodium dodecyl sulphate-SDS) with Protease Inhibitor Coctail (Sigma-Aldrich, Germany) during 2 h at 4°C. Proteins were gained by centrifugation at 12 000 g, 4°C for 20 min, and the concentrations were quantified using a BCA kit (Thermo Fisher Scientific, Slovakia) with bovine serum albumin (BSA) as a standard. Total proteins were diluted with ultra-pure water and a sample loading buffer (1: 2 solution; 10% w/v SDS, 0.02% bromophenol blue, and 25% glycerol in 0.5 M Tris-HCl, pH 6.8) and separated on 10% (for PSD95 detection) or 12% (for RhoA, RhoB detection) SDS-PAGE running gels (10 μg protein/lane) with 5% stacking gels at a constant current of 150 mA as a running parameter. The proteins were transferred to a low fluorescent polyvinylidene difluoride membrane Immobilon-FL (0.45 μm pore size; Merck-Millipore, Czech Republic) by a Mini Trans-Blot^®^ cell system (Bio-Rad, Slovakia) for 2 h in wet conditions at 300 mA. After the blocking with 4% w/v BSA (Sigma-Aldrich, Germany) for 1 h at RT, membranes were incubated overnight at +4°C with primary antibodies (Table 2). The GAPDH protein (Table 2) was used as a reference. Blots were washed three times for 10 min in Tris Buffered Saline solution with Tween^®^ and then treated with corresponding fluorescent secondary antibodies (Table 3) for 1h at RT. The Odyssey Infrared Imaging System (LI-COR, USA) was used to detect and quantify fluorescent signals in duplicate for each sample. Odyssey 2.0 analytical software (LI-COR, USA) was used to manually signpost bands of interest and quantify the density of each blot band. Results are presented as a percentage of the control group normalized to GAPDH, reference protein.

### Immunohistochemistry

For determination of excitatory synapses in the anterior dorsal CA1 and CA3 hippocampal regions in P5 WT and *Magel2^-/-^* mice, we investigated the presence of PSD95 and VGLUT2 during OT treatment (n=4 animals/group). After the perfusion with 4% AntigenFix (Diapath, Italy), brains were stored overnight in the same fixative and then sectioned on a 7000smz-2 Vibrotome (Campden Instruments LTD, UK) at a thickness of 50 μm. Sections were temporarily stored (4°C) in 24 well plates in PBS+0.01% sodium azide. For immunohistochemistry, free-floating sections were blocked with 3% donkey serum (DS), 2% BSA, 0.01% Triton X-100 in PBS. After 1 h (RT), sections were incubated with a combination of primary antibodies: anti-PSD95 and anti-VGLUT2 (Table 2) in 1% DS, 1% BSA, 0.4% Triton X-100 diluted in PBS overnight at 4°C. After washing with cold PBS (5 min/ 3times), sections were incubated with corresponding secondary antibodies (Table 3) diluted in PBS for 1.5 h at room temperature. Subsequent sections were rinsed with PBS (5 min / 3times), and 300 nM DAPI was added to stain the nuclei. After final PBS washing, sections were mounted on glass slides with Fluoromount-G (Sigma-Aldrich, Germany). PSD95- and VGLUT2-stained sections were imaged by a Nikon confocal microscope (Nikon ECLIPSE Ti-E, A1R+; Netherlands). Free-floating sections (50 μm thick) were imaged at high magnification (40x objective, numerical aperture 1.3; resolution 1024 × 1024 pixels) at 0.5 μm steps. Quantification of colocalization between VGLUT2 and PSD95 was performed using the Fiji/ImageJ plugin ComDet v0.4.1 (Bottermann et al., 2019). In brief, particles were detected from three different areas of interest in each part of the hippocampus (CA1, CA3) in both 555 and 647 channels independently with an approximate size of 2 pixels and intensity threshold of 3 SD. Colocalization was determined based on the maximum distance of 1 pixel between particles.

### Statistical Analysis

Statistical comparisons of relative gene expression and protein levels were performed using Student’s t-test. One-way ANOVA or two-way ANOVA for genotype factors and OT treatment was used when required. As a *post hoc*, the Bonferroni test or Tukey Kramer for unequal sample sizes were used. Results are expressed as the mean ± SEM. The value of p < 0.05 was considered statistically significant.

## Results

### Neurite outgrowth is impaired in *Magel2^-/-^* hippocampal cultures and compensated by OT; however, not by ATRA treatment

The morphology of neurons of hippocampal primary cultures isolated from *Magel2^-/-^* or WT neonates (the day of birth, P0) was evaluated. After dissection, the hippocampal primary cultures were cultivated for 7 days in a growing medium for the purpose of measurement of the longest neurites in immature neurons. At DIV7, cultures were treated either with or without OT (1 μM) or ATRA (10 μM) for 48 hours. Exposure of primary neurons to the positive control, ATRA, was used to examine neurite outgrowth. Statistical analysis of average neurite lengths (Figure 1) by ANOVA revealed significant differences between groups (p < 0.0001). Firstly, we compared the neurite lengths between the control, untreated WT, and *Magel2^-/-^* neurons (Figure 1a), and observed a significant 15% decrease (p < 0.001) in length of neurites (102.6±3.12μm) in the *Magel2^-/-^* neurons compared to the WT (120.9±3.37 μm). Afterwards, we examined the effects of OT or ATRA treatment in both genotypes (Figure 1a). In WT neurons, the average length of neurites significantly increased by 10% in response to OT (132.9±3.13 μm; p < 0.05) and by 30% in response to ATRA (157.5±4.02 μm; p< 0.001) incubation when compared to untreated WT neurons (120.9±3.37 μm). In the *Magel2^-/-^* neurons, we also observed a similar 13% increase of neurite outgrowth in the presence of OT (137.2±3.32 μm; p < 0.01); however, application of ATRA did not induce elongation of neurites (109.6±2.68μm). We also analyzed the number of neurons with the longest neurite expressed as a percentage of the total measured neurons. Based on the length of the neurites, we divided the neurons into 4 categories for each experimental group (Figure 1b). We found a lower percentage of neurons with the longest neurite over 150 μm in the *Magel2^-/-^* neurons (16.1%) when compared to the WT neurons (24.8%). The application of ATRA increased the number of neurons with the longest neurite in the WT neurons (55.5%); however, no effects were observed in the *Magel2^-/-^* neurons. Similarly, OT treatment increased the number of neurons with the longest neurite in the WT (35.3%) and *Magel2^-/-^* neurons (39%). The opposite effect of the *Magel2* deficiency was observed when neurites with a length up to 50 μm were evaluated. The percentage of cells with neurites up to 50 μm long was higher in the *Magel2^-/-^* neurons (14.4%) compared to the WT neurons (6.2%). By using Sholl analysis, we were able to get a detailed look at the neurite length and number of neurite branches in the neurons in a visual field (Figure 2a). We found a 27% higher sum of intersections up to 30 μm away from the nucleus in *Magel2^-/-^* (139.58±7.23) in comparison to the WT neurons (109.75±5.05). Overall, in these primary cell cultures, we observed a decrease in neurite outgrowth in *Magel2*-deficient mice versus WT mice. OT treated cell cultures increased neurite outgrowth and normalized the neurite length in *Magel2^-/-^* neurons similarly to the effect observed in WT. However, it was unexpected that ATRA treatment would have no effect on the neurite growth of *Magel2^-/-^* neurons, suggesting that the lack of *Magel2* prevents the action of ATRA.

**Fig 1:**
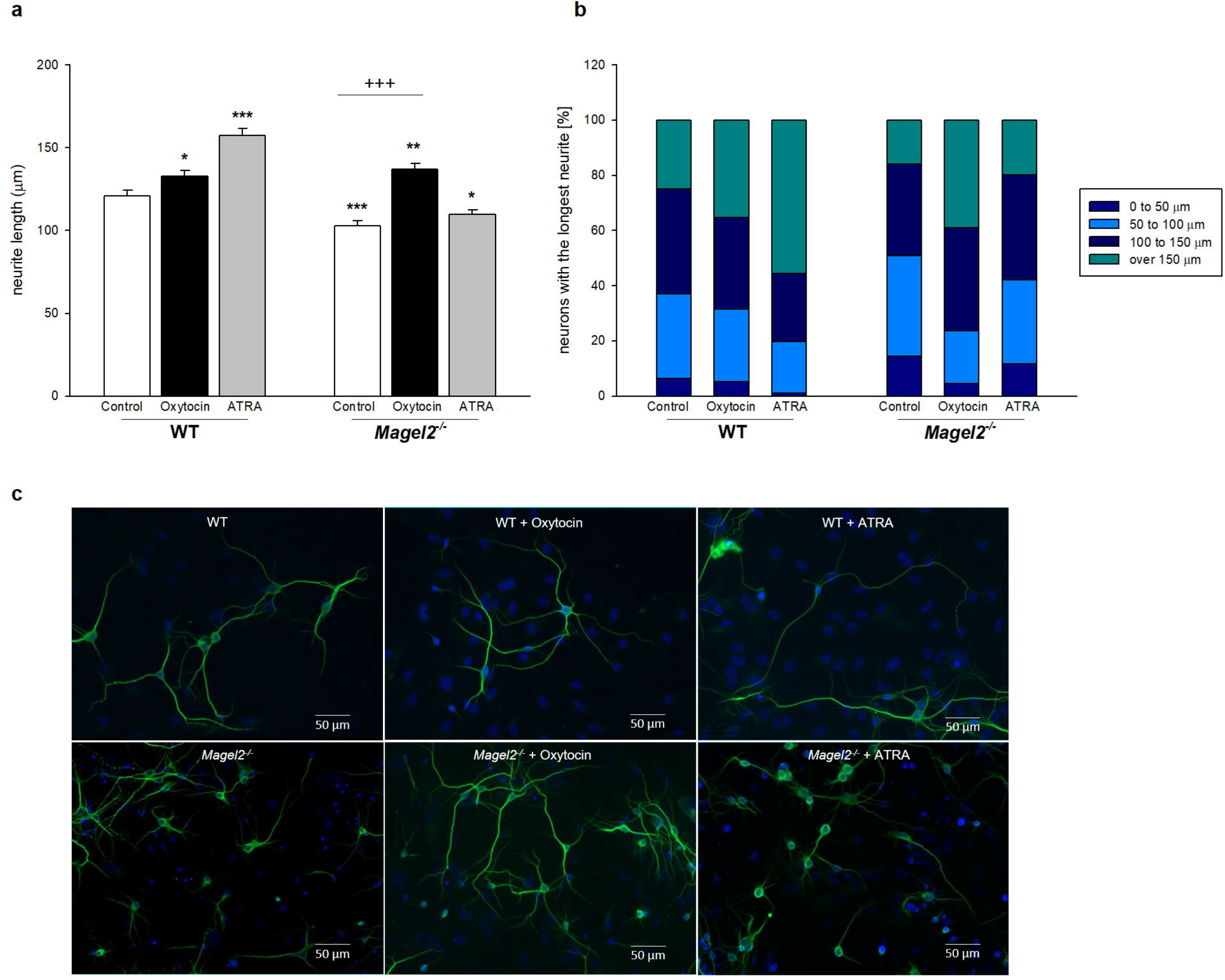
Comparative measures of neurite outgrowth of primary cultures of hippocampal neurons isolated from P0 WT or *Magel2^-/-^* mice in response to 1 μM oxytocin or 10 μM ATRA treatment at DIV9. The effect of *Magel2* deficiency, oxytocin and all-trans retinoic acid (ATRA) treatment on neurite outgrowth. Neurite length was quantified in all neurons in a visual field from the edge of the nucleus to the apical end of the neurite. The cells were plated at the density of 0.8×10^5^/ml (n =15 WT and n =7 *Magel2^-/-^* pups). Six coverslips per experimental group (n =198-317 cells) and at least 7 areas of interest per coverslip were evaluated. The average value of the longest neurite is shown. Means are represented on bars ± SEM. ANOVA revealed significant differences between groups (F_(5,1496)_ = 34.06; p < 0.0001). Tukey-Kramer *post hoc* revealed significantly different values marked with ***p < 0.001; **p < 0.01; *p < 0.05 compared to untreated control WT, **+++** p < 0.001 compared to untreated *Magel2^-/-^* **(a)**. Stacked bar graphs showing the percentage of neurons with a defined length of neurites **(b)**. Representative fluorescent microscopic images of neurons in experimental groups **(c)**. Cells were labelled for MAP2 (green) and nucleus (blue). Days *in vitro* (DIV).

**Fig 2:**
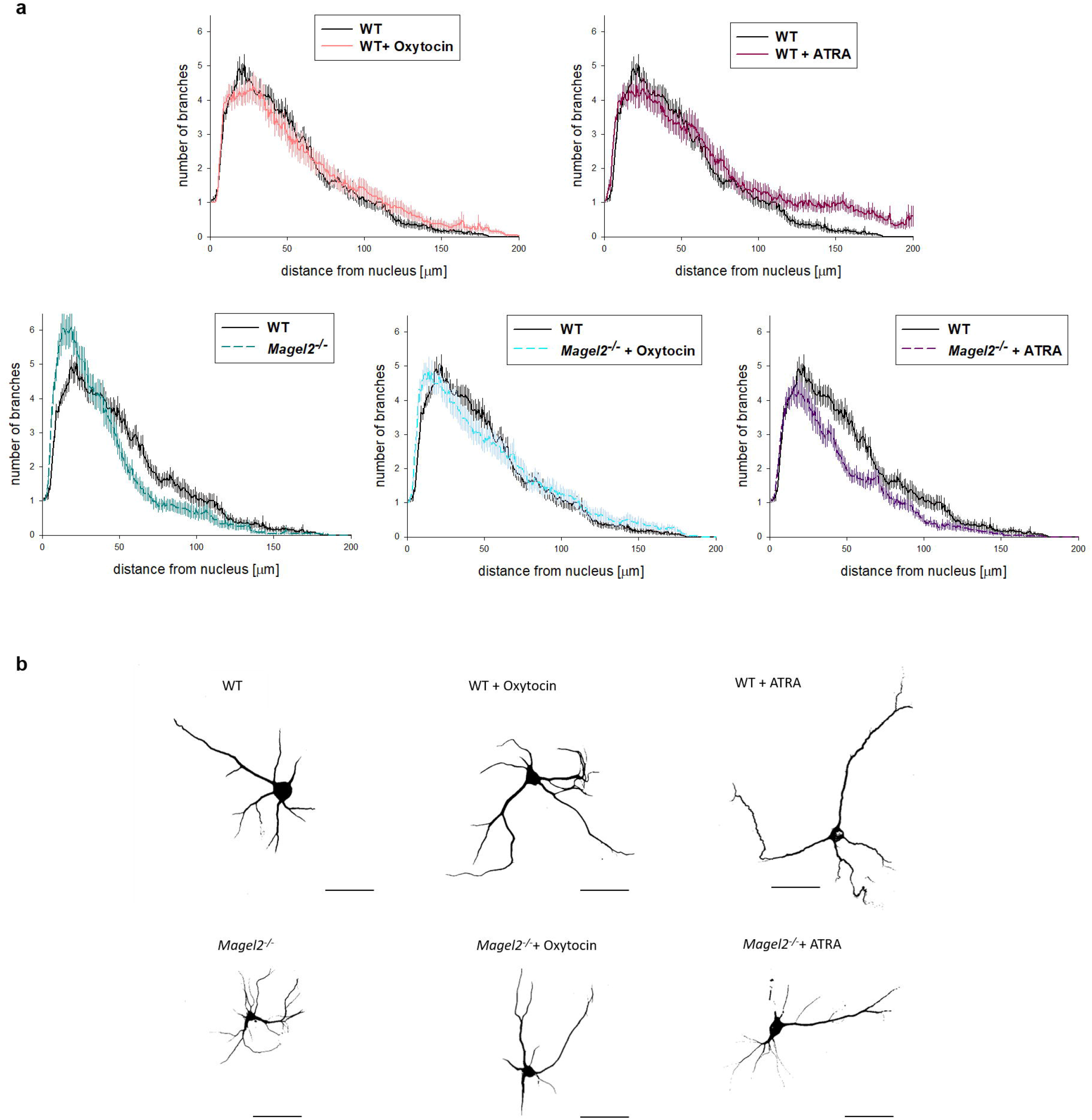
Sholl analysis of neurite arborization in primary cultures of hippocampal neurons isolated from P0 WT or *Magel2^-/-^* mice in response to 1 μM oxytocin or 10 μM ATRA treatments at DIV9. A graph showing the effect of *Magel2* deficiency, oxytocin and all-trans retinoic acid (ATRA) treatment. The number of neurite intersections for concentric circles with various distances away from the nucleus was calculated using Fiji/ImageJ **(a)**. Representative binary images of neurons **(b)** for all experimental groups are shown (n = 40 cells per group). The cells were plated at the density of 0.8×10^5^/ml (n =15 WT and n =7 *Magel2^-/-^* pups). Scale bar represents 50 μm. Days *in vitro* (DIV).

### The quantity of neurite outgrowth-associated transcripts is altered during development of the hippocampus in *Magel2^-/-^* mice compared with WT controls

In order to further study the deficit in neurite outgrowth in *Magel2^-/-^* mice versus the WT genotype, we looked at differences in mRNA transcripts or protein levels of specific neurite outgrowth associated proteins (Figure 3) at two different developmental stages: in five day-old pups (P5) and in 30 day-old mice (P30) in the hippocampal tissue. We observed several changes in *Magel2*-deficient mice. Indeed, at P5, the mRNA levels of *Growth associated protein 43-Gαp43* (0.73±0.09; p < 0.001) and *p21 activated kinase 1* - *Pak1* (0.74±0.12; p < 0.01) were significantly reduced compared to the WT mice (Figure 3a). The quantity of *Map2* transcripts was not different, however, between genotypes. Furthermore, we quantified expression of *Ras-related C3 botulinum toxin substrate* - *Rac1*, chosen for its highly-recognized roles within Rho GTPase signaling, which is important in regulating the cytoskeleton involved in axonal outgrowth (Cao, Deng & Pocock 2019). Because ATRA did not have any effect on neurite outgrowth in *Magel2*-deficient mice, we also evaluated GTP-dependent upstream kinase *T-cell lymphoma invasion and metastasis 1* - *Tiam1* and downstream kinase *LIM domain kinase 1* - *Limk1*, both enzymes associated with rearranging of the cytoskeleton and interact with the retinoid receptor function (Ishaq et al., 2011). At P5, no difference in *Rac1* expression was found; however, we observed lower mRNA levels of *Tiam1* (0.43±0.10; p < 0.001) and *Limk1* (0.82±0.10; p < 0.05) compared to the WT (Figure 3b). Tiam1 and Limk1 are involved in signaling cascades under the control of small Rho GTPases and the Ras homolog family members A and B (RhoA and RhoB) as well. We then studied the RhoA and RhoB transcripts and proteins in *Magel2^-/-^* mice (Figure 4). At P5, expression of *RhoB* increased at the transcript (1.60±0.19; p < 0.01) and protein (171.20±25.89; p < 0.05) level compared to the WT; however, such changes were not observed for the expression of *RhoA* (Figure 4a, 4b).

**Fig 3:**
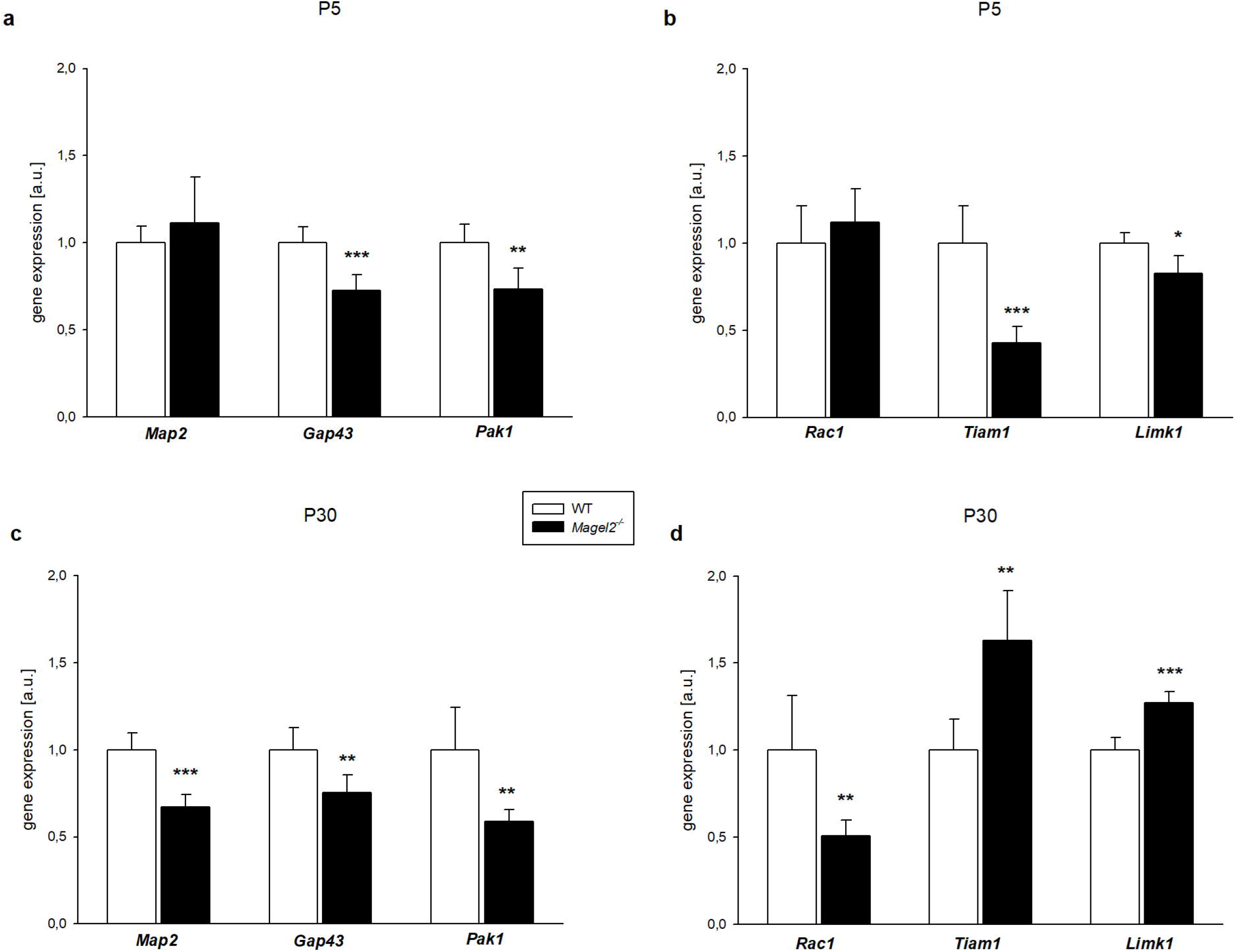
Changes in the hippocampal neurite outgrowth-associated transcript levels at P5 (a, b) and P30 (c, d) WT or *Magel2^-/-^* mice. The graphs show relative mRNA expression levels normalized to *Gapdh* transcript. qRT-PCR was calculated by the 2^-ΔΔCt^ Livak method (Livak & Schmittgen, 2001). Neurite outgrowth-associated transcripts at P5 **(a)** and P30 **(c)**; small GTPases and kinases at P5 **(b)** and P30 **(d)**. Means are represented on bars ± SEM (n = 4-6/group). At P5, the mRNA levels of *Gap43* (***p < 0.001, df=9, T=5.01), *Pak1* (**p < 0.01, df=9, T=3.83), *Tiam1* (***p < 0.001, dt=9, T=5.94) and *Limk1* (*p < 0.05, df=7, T=3.01) were significantly reduced in *Magel2^-/-^* mice compared to the WT mice. At P30, the mRNA levels of Gap43 (**p < 0.01, df=10, T=3.71), Pak1 (**p < 0.01, df=9, T=3.97), Map2 (***p < 0.001, df=10, T=6.62) and Rac1 (**p < 0.01, df=10, T=3.72) were significantly lower *in Magel2^-/-^* mice compared to the WT mice. At P30, Tiam1 (**p < 0.01, df=10, T=4.55) and Limk1 (***p < 0.001, df=10, T=6.85) gene expression levels increased in *Magel2^-/-^* mice compared to the WT mice. *Microtubule associated protein 2* - *Map2; Growth associated protein 43* - *Gap43; p21 activated kinase 1* - *Pak1; Ras-related C3 botulinum toxin substrate 1* - *Rac1; T-Lymphoma invasion and metastasis-inducing protein 1* - *Tiam1; LIM domain kinase 1* - *Limk1*.

**Fig 4:**
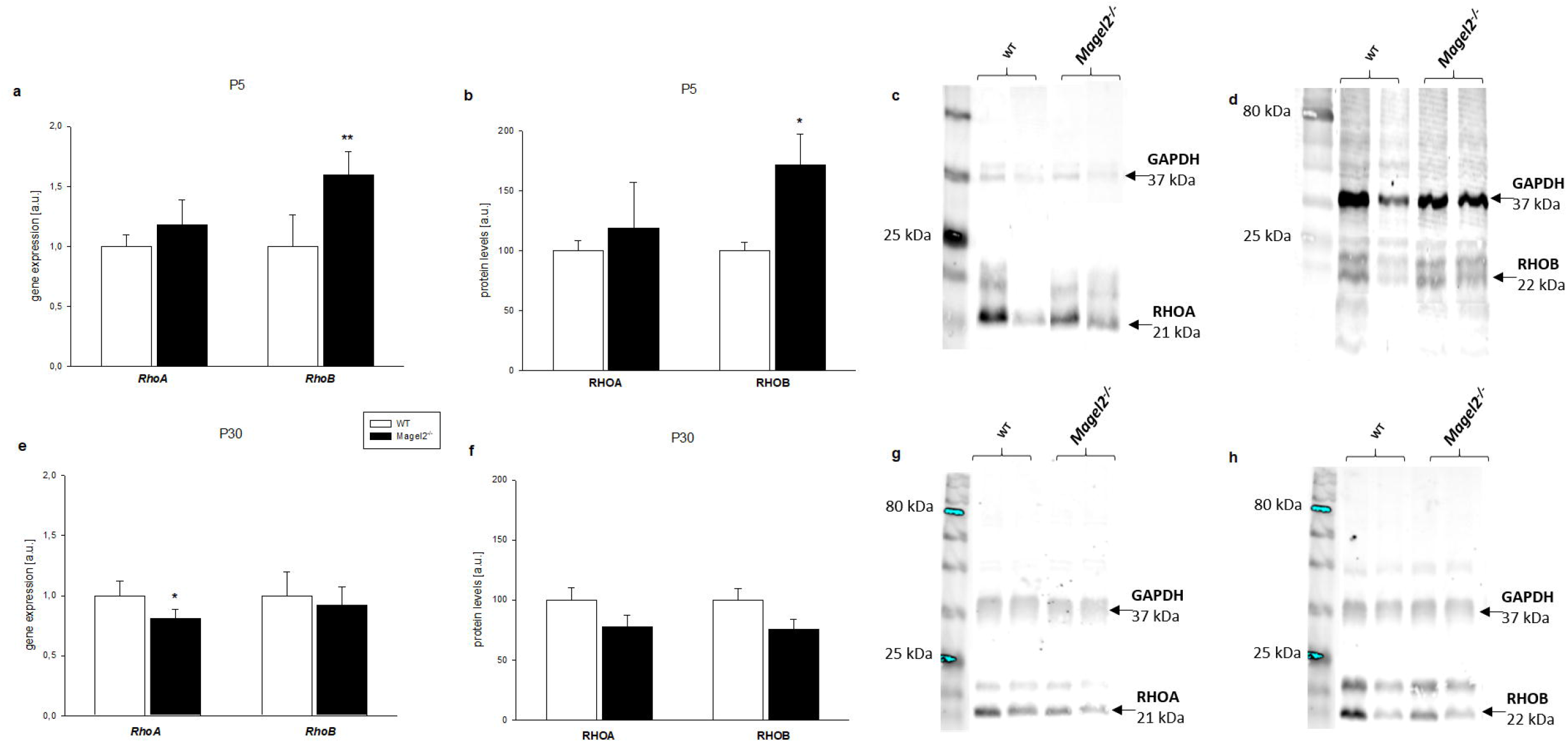
Changes in the hippocampal transcript and protein levels of RhoA and RhoB at P5 (a-d) and P30 (e-h) WT or *Magel2^-/-^* mice. The graphs show relative mRNA expression levels **(a, e)** normalized to *Gapdh* transcript and protein levels **(b, f)** normalized to GAPDH protein. qRT-PCR was calculated by the 2^-ΔΔCt^ Livak method (Livak & Schmittgen, 2001). Representative set of Western Blot Analysis results from RHOA **(c, g)** and RHOB **(d, h)**. Means are represented on bars ± SEM (n = 4-6/group). At P5, *RhoB* (RHOB) mRNA (**p < 0.01, df=9, T=4.34) and protein (*p < 0.05, df=6, T=2.66) levels were significantly increased in *Magel2^-/-^* mice compared to the WT mice. At P30, the mRNA level of *RhoA* (*p < 0.05, df=10, T=3.25) was significantly lower in *Magel2^-/-^* mice compared to the WT mice. *Ras homolog family member – RhoA and RhoB*.

At P30, the same significant level of reduction for *Gap43* (0.75±0.10; p < 0.01) and for *Pak1* (0.59±0.07; p < 0.01) was observed along with lower mRNA levels (0.67±0.07; p < 0.001) of *Map2* in *Magel2^-/-^* mice (Figure 3c); mRNA levels of *Rac1* were also significantly lower (0.51±0.09; p < 0.01) in *Magel2^-/-^* mice when compared to the WT mice. Conversely, *Tiam1* (1.63±0.29; p < 0.01) and *Limk1* (1.27±0.06; p < 0.001) gene expression levels increased in *Magel2^-/-^* mice (Figure 3d). At P30, we observed a significant decrease of the mRNA level of *RhoA* (0.81±0.08; p < 0.05) in *Magel2*-deficient mice and a similar trend towards a decrease in the protein levels of RHOA (77.87±9.43; ns) in *Magel2^-/-^* mice was found; RhoB was expressed at the same level in both genotypes (Figure 4e, 4f). In conclusion, in P5 immature neurons, we observed a significant reduction in the quantity of several transcripts for factors involved in neurite outgrowth in *Magel2^-/-^* mice. At P30, in mature neurons, most of the reduced markers observed at P5 had a normal or even higher level of expression.

### The quantity of presynaptic transcripts is reduced at P5 and normalized or increased at P30 in *Magel2^-/-^* mice compared to the WT mice

The synapse-associated molecules (Figure 5) were also examined. At days P5 and P30, we investigated two splicing variants of cell-adhesion molecules, neurexins (Nrxns), and two synaptic vesicle proteins: Synapsin I and Synaptophysin; all of which are known to be present in the “presynaptic compartments”. At P5, the quantity of *Nrxn1α* and *Nrxn2β* transcripts was significantly lower (0.87±0.05; p < 0.05 and 0.67±0.13; p < 0.01, respectively) in *Magel2*-deficient mice compared to the WT mice. No change was observed for mRNA levels of *Nrxn1β* and *Nrxn2α* (Figure 5a). We also observed a decrease of *Synapsin I* (0.86±0.05; p < 0.05) and *Synaptophysin* (0.87±0.09; p < 0.05) transcripts in *Magel2^-/-^* mice (Figure 5b). At P30, we found that in *Magel2^-/-^* mice, the quantity of *Nrxn1β* (1.16±0.02; p < 0.05), *Nrxn2α* (1.19±0.07; p < 0.001) and *Nrxn2β* (1.23±0.09; p < 0.01) transcripts was significantly higher compared to the WT mice (Figure 5c). No differences were found in mRNA expression of *Synapsin I* and *Synaptophysin* at P30 (Figure 5d). Altogether, these data suggest a significant decrease in the quantity of presynaptic markers at P5 in *Magel2*-deficient mice compared to the WT mice; this deficit appears normalized at P30 with even a slight increase.

**Fig 5:**
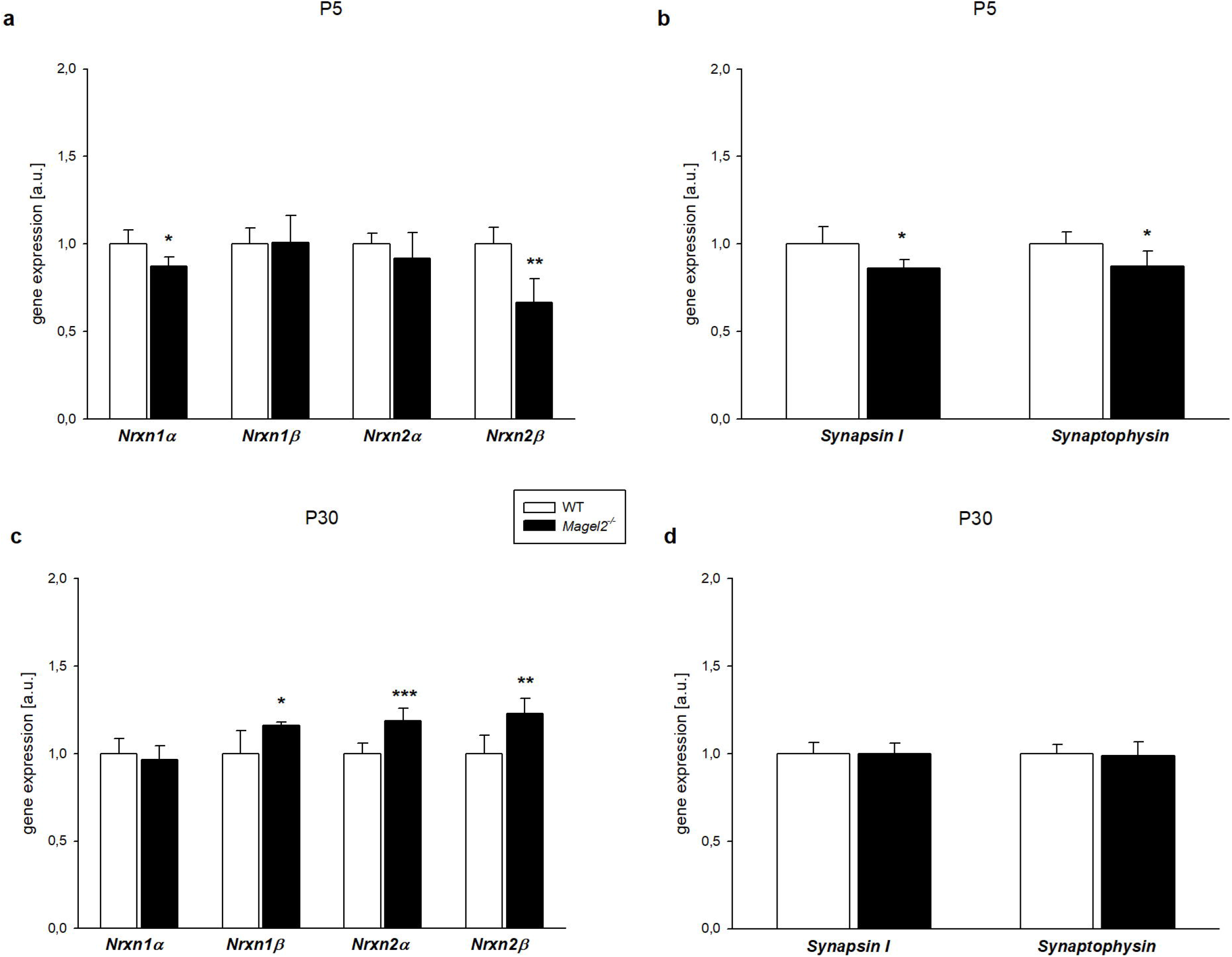
Changes in the hippocampal presynaptic transcript levels at P5 (a, b) and P30 (c, d) WT or *Magel2^-/-^* mice. The graphs show relative mRNA expression levels normalized to *Gapdh* transcript. qRT-PCR was calculated by the 2^-ΔΔCt^ Livak method (Livak & Schmittgen, 2001). Presynaptic celladhesion molecules at P5 **(a)** and P30 **(c)**; synaptic vesicle molecules at P5 **(b)** and P30 **(d)**. Means are represented on bars ± SEM (n = 4-6/group). At P5, the mRNA levels of *Nrxn1α* (*p < 0.05, df=8, T=2.99), *Nrxn2β* (**p< 0.01, df=9, T=4.67), *Synapsin I* (*p < 0.05, df=9, T=3.03) and *Synaptophysin* (*p < 0.05, df=9, T=2.44) were significantly reduced in *Magel2^-/-^* mice compared to the WT mice. At P30, *Nrxn1β* (*p < 0.05, df=10, T=2.99), *Nrxn2α* (***p < 0.001, df=10, T=5.13) and *Nrxn2β* (**p < 0.01, df=10, T=4.09) gene expression levels increased in *Magel2^-/-^* mice compared to the WT mice. *Neurexins* - *Nrxns*.

### Shank and Neuroligin postsynaptic transcripts are increased in *Magel2^-/-^* mice compared to the WT mice at P30 but not at P5

Additionally, we also examined whether *Magel2* deficiency plays a role in the regulation of postsynaptic transcripts and proteins in the hippocampal tissue (Figure 6). We measured levels of scaffolding and cell-adhesion molecules involved in the balance between excitatory and inhibitory synapses (Prange et al., 2004; Varoqueaux et al., 2006). At P5, no differences were found in transcripts of *“SH3 and multiple ankyrin repeat domains” (Shank1-3*) and *Neuroligins (Nlgn1-2*) genes in *Magel2^-/-^* mice compared to the WT mice (Figure 6a, 6b). *Nlgn3* transcript was the only slightly increased postsynaptic gene in *Magel2^-/-^* mice at P5 (1.15±0.13; p < 0.05) (Figure 6b). In contrast to early development, several cases of genes coding for postsynaptic transcripts were upregulated at P30 (Figure 6c, 6d). Indeed, we found that in *Magel2^-/-^* mice, the *Shank1* (1.22±0.11; p < 0.001), *Shank3* (1.17±0.07; p < 0.01), *Nlgn2* (1.37±0.07; p < 0.001) and *Nlgn3* (1.17±0.10; p < 0.05) transcripts were significantly higher compared to the WT mice. Conversely, *Nlgn1* transcript was decreased (0.82±0.05; p < 0.01) in *Magel2^-/-^* mice compared to the WT mice. The expression of *Postsynaptic density protein 95* (*Psd95*) was also regulated differently in P5 and P30 (Figure 7). We found significantly lower (0.77±0.05; p < 0.001) gene expression and the same trend for the protein levels of PSD95 (85.55±15.47; ns) in P5 *Magel2*-deficient mice (Figure 7a, 7b). However, the trend for *Psd95* changes at P30 was the opposite. Statistical analysis revealed no change in gene expression, but significantly higher (138.90±15.95; p < 0.05) protein levels of PSD95 at P30 (Figure 7e). In conclusion, at P5, a quantity of postsynaptic transcripts and proteins revealed no change globally, yet only a small increase of *Nlgn3* and decrease of PSD95. At P30, we found an increase of *Shank* and *Nlgn* transcripts, but a decrease of *Nlgn1* transcript in *Magel2^-/-^* mice compared to the WT mice and an increase of PSD95.

**Fig 6:**
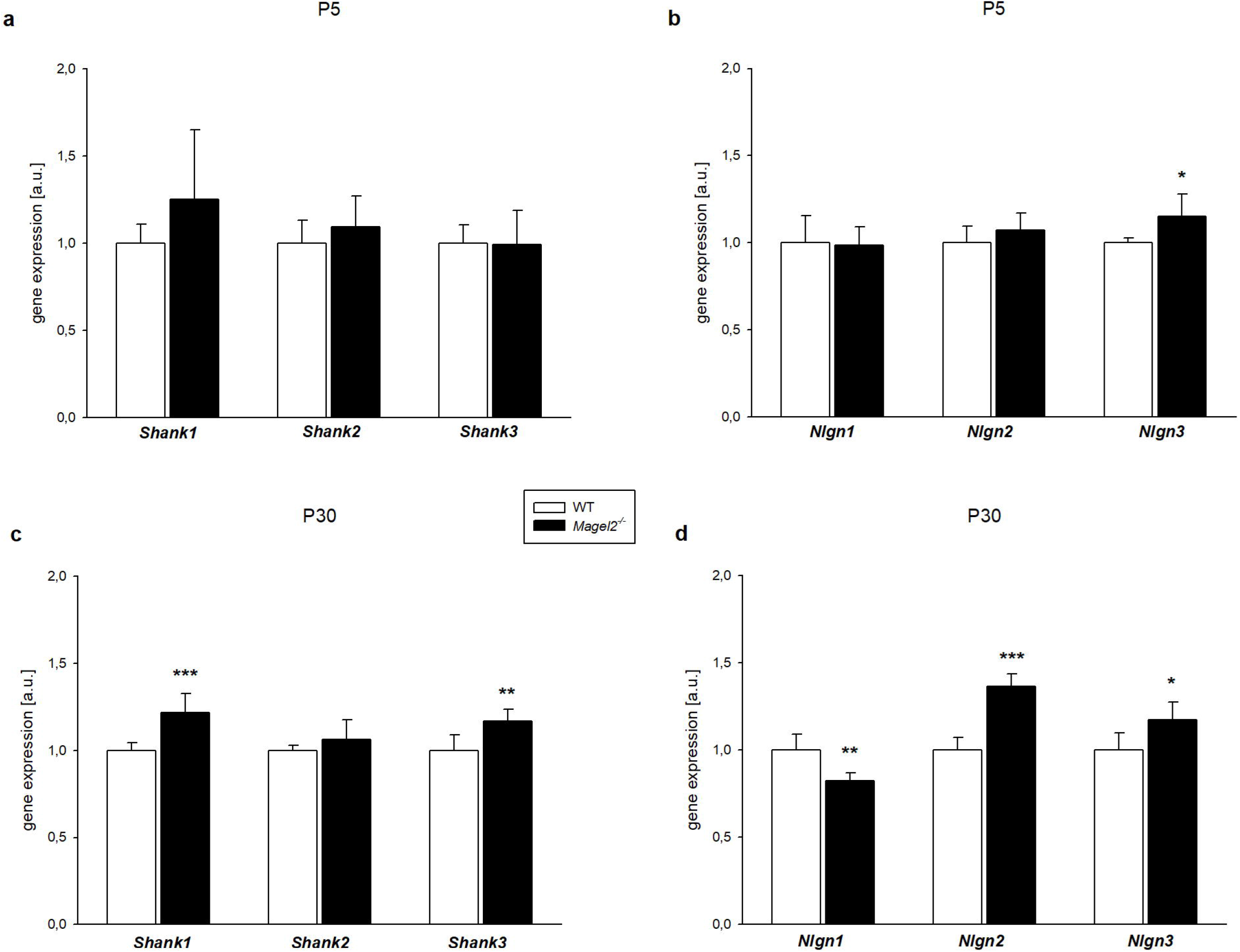
Changes in the hippocampal postsynaptic transcript levels at P5 (a, b) and P30 (c, d) WT or *Magel2^-/-^* mice. The graphs show relative mRNA expression levels normalized to *Gapdh* transcript. qRT-PCR was calculated by the 2^-ΔΔCt^ Livak method (Livak & Schmittgen, 2001). Scaffolding proteins at P5 **(a)** and P30 **(c)**; postsynaptic cell adhesion molecules at P5 **(b)** and P30 **(d)**. Means are represented on bars ± SEM (n = 4-6/group). At P5, the mRNA level of *Nlgn3* (*p < 0.05, df=9, T=2.57) was increased in *Magel2^-/-^* mice compared to the WT mice. At P30, the mRNA levels of *Shank1* (*** p < 0.001, df=10, T=4.60), *Shank3* (** p < 0.01, df=10, T=3.67), *Nlgn2* (*** p < 0.001, df=10, T=9.00) and *Nlgn3* (* p < 0.05, df=10, T=3.05) were significantly higher *in Magel2^-/-^* mice compared to the WT mice. At P30, *Nlgn1* (**p < 0.01, df=10, T=4.38) gene expression level decreased in *Magel2^-/-^* mice compared to the WT mice. *SH3 and multiple ankyrin repeat domains* - *Shanks; Neuroligins* – *Nlgns*.

**Fig 7:**
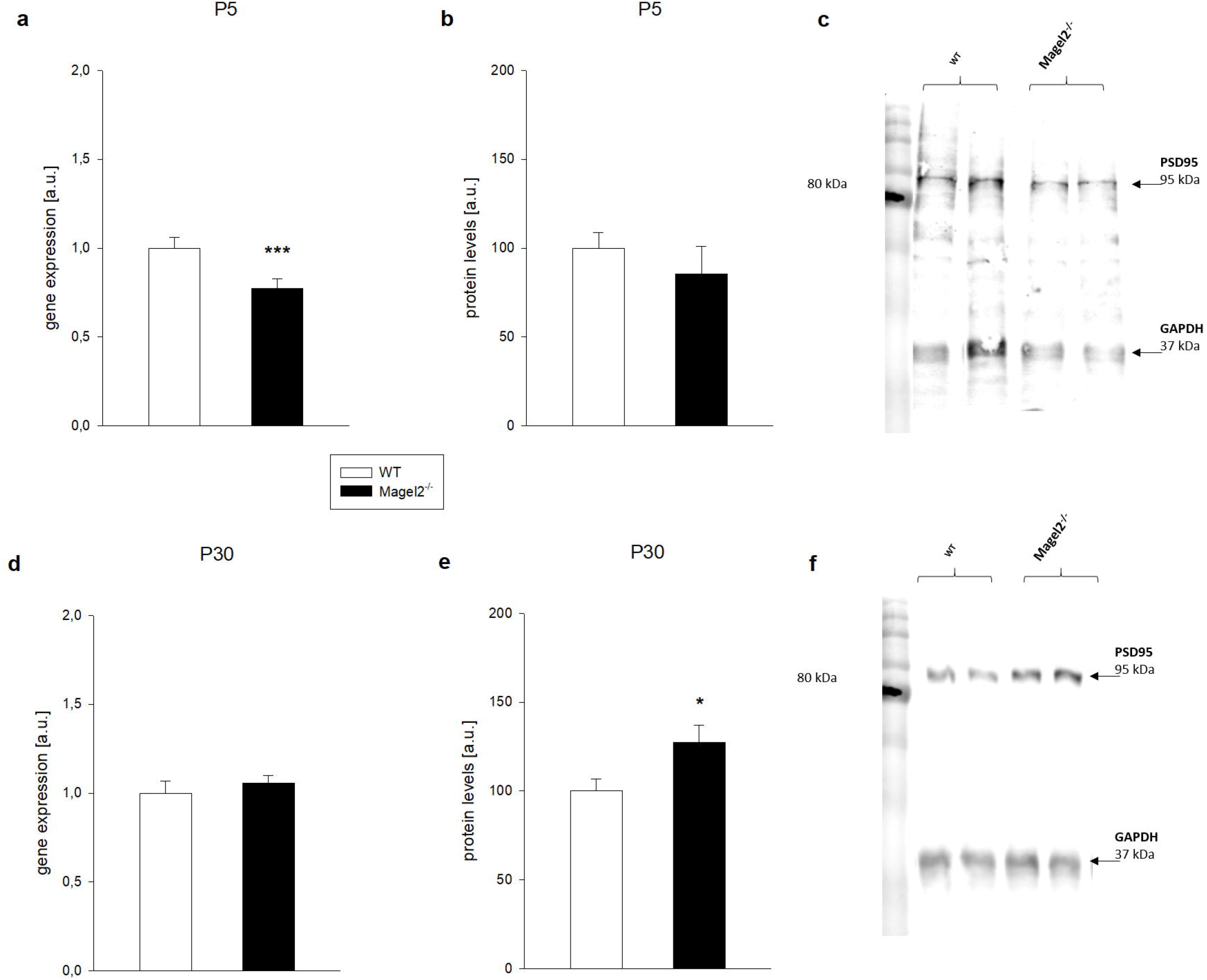
Changes in the hippocampal transcript and protein levels of Psd95 at P5 (a-c) and P30 (d-f) WT or *Magel2^-/-^* mice. The graphs show relative mRNA expression levels **(a, d)** normalized to *Gapdh* transcript and protein levels **(b, e)** normalized to GAPDH protein. qRT-PCR was calculated by the 2^-ΔΔCt^ Livak method (Livak & Schmittgen, 2001). Representative set of Western Blot Analysis results from PSD95 at P5 **(c)** and P30 **(f)**. Means are represented on bars ± SEM (n = 4-6/group). At P5, the mRNA level of *Psd95* (*** p < 0.001, df=8, T=6.15) was significantly reduced in *Magel2^-/-^* mice compared to the WT mice. At P30, PSD95 (* p < 0.05, df=9, T=2.29) protein level increased in *Magel2^-/-^* mice compared to the WT mice. *Postsynaptic density protein 95 - Psd95*.

### OT stimulates the expression of presynaptic and postsynaptic transcripts in P5 *Magel2^-/-^* mice compared to the WT mice

We also investigated the effect of OT treatment (2μg/pup, s.c.) administered at P2 and P3 on the levels of presynaptic and postsynaptic transcripts (values and statistics are shown in Table 5). At P5, following OT treatment, we observed significantly higher levels of *Nrxn1β* (p < 0.001), *Nrxn2α* (p < 0.01), *Nlgn1* (p < 0.01) and *Nlgn2* (p < 0.001) transcripts in *Magel2*-deficient mice compared to the WT mice. The quantity of those transcripts was unchanged in *Magel2^-/-^* mice compared with the WT mice. Furthermore, we observed a significant decrease in the quantity of *Psd95* (p < 0.001), *Nrxn1α* (p < 0.05) and *Nrxn2β* (p < 0.01) in *Magel2*-deficient mice compared with the WT mice (Table 5), and OT treatment normalized the quantity of those transcripts. Globally, OT treatment enhanced the expression of all the previous studied transcripts at P5 in *Magel2*-deficient mice except for *Nlgn3*.

**Table 5:** Changes in the hippocampal presynaptic and postsynaptic transcript levels in response to oxytocin treatment at P5 WT or *Magel2^-/-^* mice. Values show relative mRNA levels normalized to *Gapdh* transcript. qRT-PCR was calculated by the 2^-ΔΔCt^ Livak method (51). *Neurexins* – *Nrxns; Neuroligins* – *Nlgns; Postsynaptic density protein 95* - *Psd95*. Represented are means ± SEM (n = 4-6/group). Significantly different values are marked with ***p<0.001; **p<0.01 and *p<0.05 compared to WT.

|  | WT | *Magel2*<sup>-/-</sup> |  | *Magel2*<sup>-/-</sup> + Oxytocin |  |
| --- | --- | --- | --- | --- | --- |
| *gene* | mRNA levels | mRNA levels | statistics (df; T) | mRNA levels | statistics (df; T) |
| *Nrxn1α* | 1±0.08 | **0.87±0.05\*** | **8; 3.00** | 1.10±0.10 | 11; 1.08 |
| *Nrxn1β* | 1±0.09 | 1.01±0.15 | 9; 0.11 | **1.39±0.11\*\*\*** | 11; 6.74 |
| *Nrxn2α* | 1±0.06 | 0.92±0.14 | 9; 1.17 | **1.19±0.10\*\*** | 11; 4.02 |
| *Nrxn2β* | 1±0.09 | **0.67±0.13\*\*** | **9; 4.67** | 1.01±0.11 | 11; 0.09 |
| *Nlgn1* | 1±0.15 | 0.99±0.10 | 9; 0.16 | **1.29±0.13\*\*** | 11; 3.57 |
| *Nlgn2* | 1±0.09 | 1.07±0.10 | 9; 1.23 | **1.52±0.11\*\*\*** | 11; 8.91 |
| *Nlgn3* | 1±0.03 | **1.15±0.13\*** | **9; 2.57** | 1.05±0.17 | 11; 0.75 |
| *Psd95* | 1±0.06 | **0.77±0.05\*\*\*** | **8; 6.15** | 1.10±0.19 | 10; 1.31 |

### A lower immunofluorescence signal of excitatory presynaptic VGLUT2 and postsynaptic PSD95 markers in *Magel2^-/-^* compared to WT primary hippocampal neurons

For the analysis of excitatory synaptic proteins in primary hippocampal neurons, triple staining for PSD95, VGLUT2, and MAP2 was performed using specific antibodies. The total quantity of fluorescence of the VGLUT2 normalized on MAP2 (VGLUT2/MAP2; Figure 8a) decreased by 21% in primary hippocampal neurons isolated from the *Magel2^-/-^* (3.06±0.11; p < 0.01) mice compared to the WT (3.87±0.29) mice. Similarly, the signal intensity of PSD95/MAP2 significantly decreased by 37% in *Magel2^-/-^* (0.86±0.02; p < 0.001) compared with WT (1.37±0.08) neurons (Figure 8b). These results suggest a decrease of markers of glutamatergic synapses in *Magel2^-/-^* hippocampal cultures compared with the WT.

**Fig 8:**
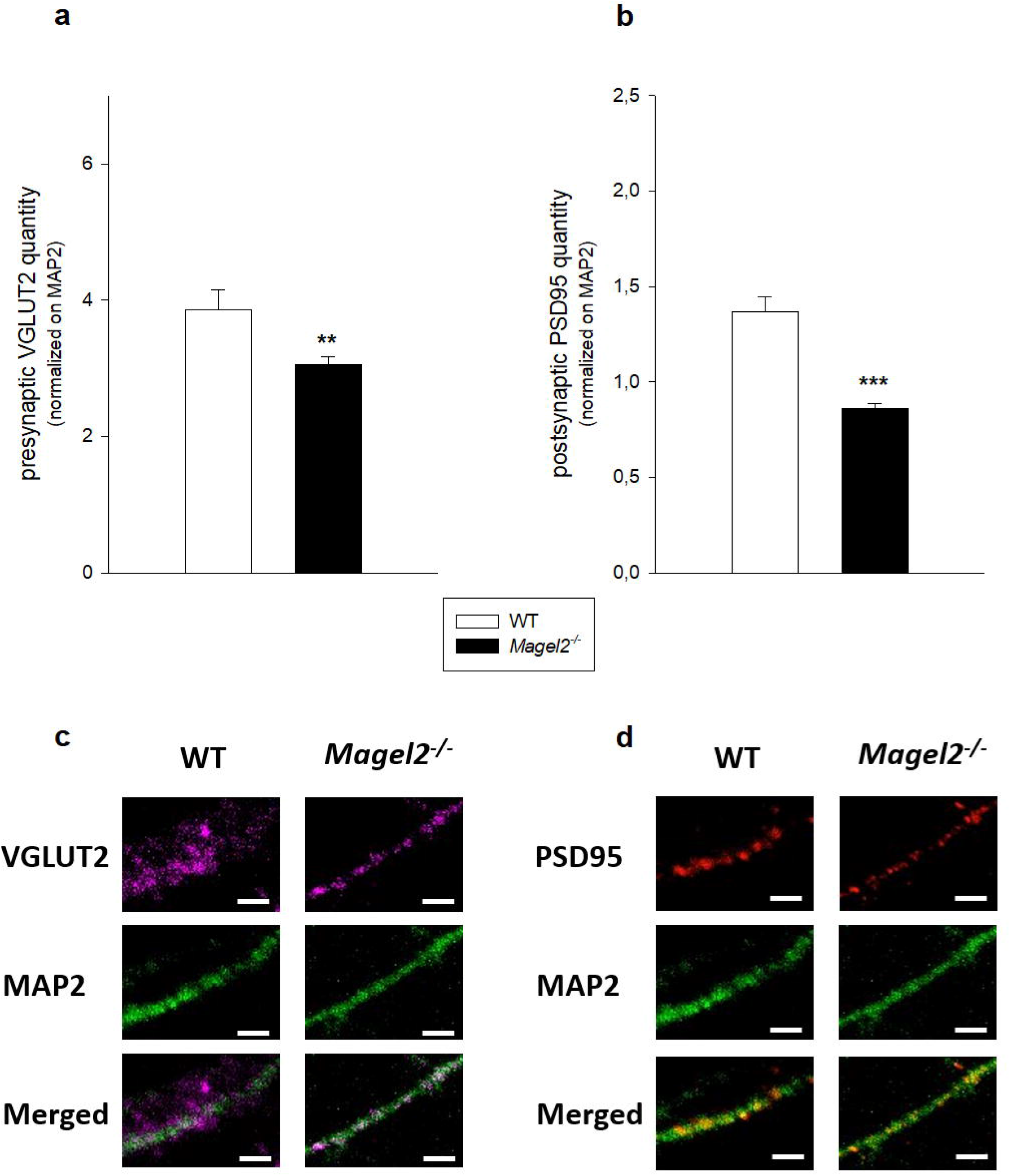
Quantitative assessment of excitatory presynaptic VGLUT2 and postsynaptic PSD95 markers in primary cultures of hippocampal neurons isolated from P0 WT or *Magel2^-/-^* mice at DIV9. The graphs show the effect of *Magel2* deficiency on excitatory presynaptic VGLUT2 **(a)** and postsynaptic PSD95 **(b)** markers. Representative fluorescent microscopic images of neurites as they were used for quantification **(c, d)**. Quantitative analysis of the fluorescent staining of presynaptic VGLUT2-positive and PSD95-positive compartments juxtaposed to MAP2-positive neurites, expressed in total quantity of fluorescence. Three individual dendritic segments (10 μm each) per neuron at least 10 μm from the nucleus were randomly chosen. The scale bar represents 2 μm. The cells were plated at the density of 0.8×10^5^/ml (n =9 WT and n =9 *Magel2^-/-^* pups). Means are represented on bars ± SEM (n = 110-150 cells). Student’s t-test revealed a decrease of the total quantity of fluorescence of the VGLUT2 (**p < 0.01, df=224, T=2.78) in *Magel2^-/-^* compared to the WT neurons **(a)**. Student’s t-test revealed a decrease of the total quantity of fluorescence of the PSD95 (***p < 0.001, df=219, T=6.39) in *Magel2^-/-^* compared to the WT neurons **(b)**. Postsynaptic density protein 95 (PSD95); Vesicular Glutamate Transporter 2 (VGLUT2); Microtubule Associated Protein 2 (MAP2).

### At P5, *Magel2^-/-^* mice manifest a lower immunofluorescence signal for PSD95/VGLUT2 colocalization in the hippocampal CA1 and CA3 regions compared with the WT mice, and OT treatment decreases the signal in WT mice

In order to gain more insight into the excitatory synaptic connections in the hippocampus, we quantified the anterior dorsal areas of CA1 and CA3 regions for the PSD95/VGLUT2 colocalization in the tissue isolated from P5 mice (Figure 9). A significant 44% decrease of the colocalization in the hippocampal area CA1 in *Magel2^-/-^* (27.68±4.64; p < 0.05) mice compared to the WT (49.36±6.69) mice was found (Figure 9a). Surprisingly, OT treatment decreased the quantity of colocalized signal by 61% in the hippocampal area CA1 in WT mice (19.21±2.57; p < 0.001) without any additional effect in *Magel2*-deficient mice (28.19±4.12; p < 0.05). A significant 29% decrease in the colocalization of PSD95/VGLUT2 in the hippocampal area CA3 in *Magel2*-deficient mice (68.99±7.45; p < 0.05) compared to the WT mice (97.55±7.31) was found (Figure 9b). Administration of OT decreased the quantity of the colocalized signal by 32% in the WT mice only (66.30±4.85; p < 0.05). Therefore, we confirmed a decrease in markers of glutamatergic synapses in brain tissue in the CA1 and CA3 hippocampal regions of *Magel2^-/-^* at P5 that is not rescued after OT treatment at P2-P3. Unexpectedly, we observed a strong and significant reduction of markers of glutamatergic synapses in WT pups treated with OT.

**Fig 9:**
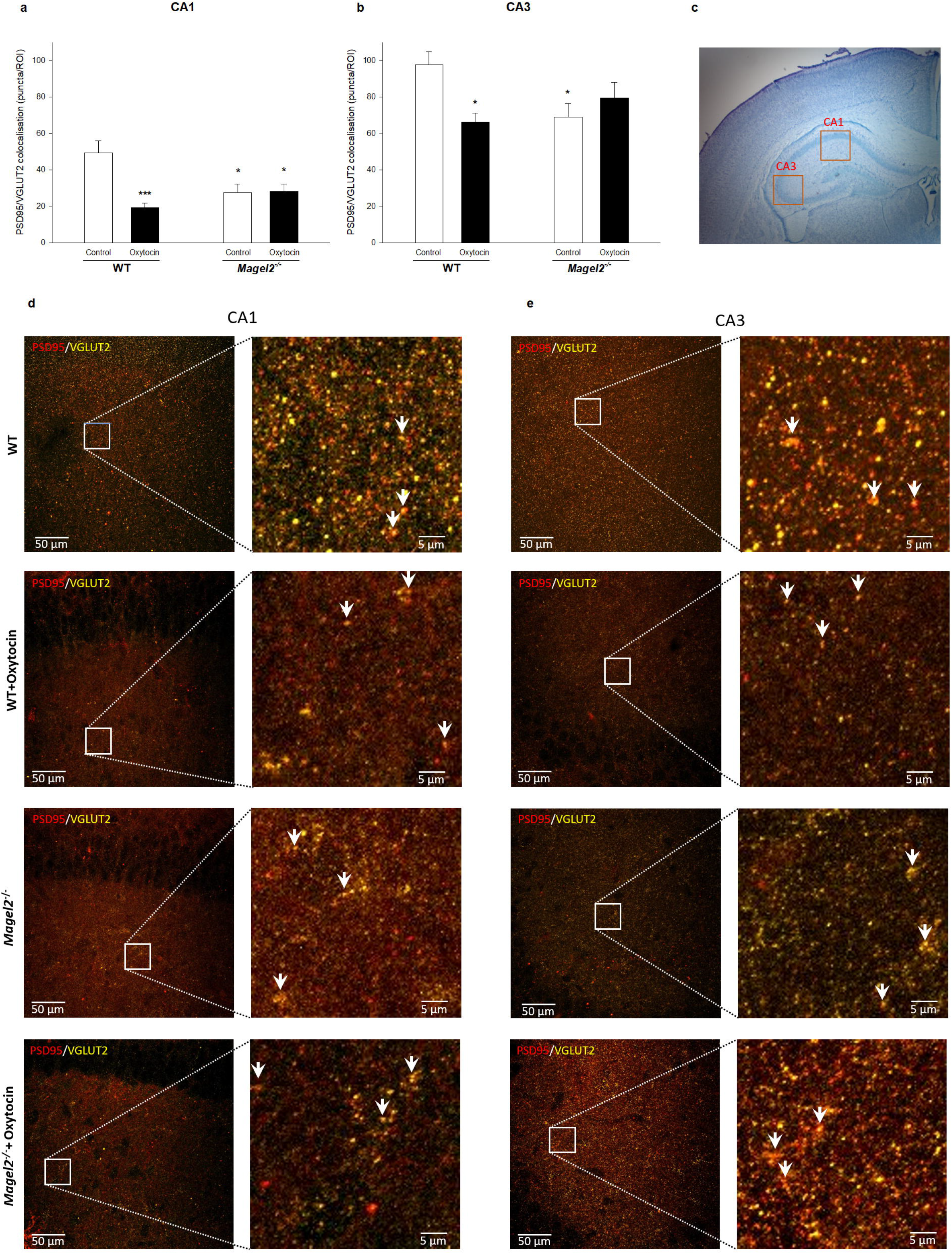
Quantification of colocalization between VGLUT2 and PSD95 in the hippocampal areas CA1 and CA3 WT or *Magel2^-/-^* mice. The graphs show the effect of *Magel2* deficiency and oxytocin treatment on colocalization of PSD95 and VGLUT2 in the hippocampal areas CA1 **(a)** and CA3 **(b)**. Representative image of chosen regions **(c)** and microscopic images showing immunoreactivity to PSD95/VGLUT2 **(d,e)**. Quantification of colocalization between PSD95 and VGLUT2 was performed using the Fiji/ImageJ plugin ComDet v0.4.1. Particles were detected from three different regions of interest (ROI) in each part of the hippocampus (CA1, CA3) in both 555 and 647 channels independently with an approximate size of 2 pixels and intensity threshold of 3 SD (n=4 animals/group, 10 sections of bilateral hippocampus/group, 3 ROI/ each part of the hippocampus). Colocalization was determined based on maximum distance of 1 pixel between particles. Means are represented on bars ± SEM. In the CA1 hippocampal area **(a)**, two-way ANOVA revealed significant differences for the factor treatment (F_(1,70)_ = 8.294, p < 0.01), as well as for the interaction of factors genotype and oxytocin treatment (F_(1,70)_ = 8.885, p < 0.01). The Bonferroni *post hoc* test revealed a significant decrease (* p < 0.05) of the PSD95/VGLUT2 colocalization in *Magel2^-/-^* mice compared to the WT mice. Oxytocin treatment decreased the quantity of colocalized signal in both the WT (***p < 0.001) and *Magel2* deficient mice (*p < 0.05). In the CA3 hippocampal area **(b)**, two-way ANOVA revealed a significant difference in the interaction of factors genotype and oxytocin treatment (F_(1,76)_ = 8.420, p < 0.01). The Bonferroni *post hoc* test revealed a significant decrease in the PSD95/VGLUT2 colocalization CA3 in *Magel2* deficient mice (*p < 0.05) compared to the WT mice. Administration of oxytocin to the WT mice decreased the quantity of colocalized PSD95/VGLUT2 signal (*p < 0.05). Postsynaptic Density Protein 95 (PSD95); Vesicular Glutamate Transporter 2 (VGLUT2).

### The quantity of glutamic acid decarboxylase transcripts and proteins are increased in the hippocampus of *Magel2^-/-^* mice compared to the WT mice

In order to achieve a global view of the GABAergic activity, we investigated the quantity of two isoforms of glutamic acid decarboxylase (GAD65 and GAD67), the enzyme that catalyzes the formation of γ-aminobutyric acid (GABA) from glutamic acid at both developmental stages. GAD67 is constitutively active and produces more than 90% of GABA; GAD65 is transiently activated and augments GABA levels for rapid modulation of inhibitory transmission. Thus, we examined whether inhibitory transmission is affected in *Magel2^-/-^* mice. At P5 (Figure 10a and 10b), we found a significant increase of *Gad65* (1.38±0.05; p < 0.01) and *Gad67* (1.53±0.24; p<0.001) transcripts and a non-significant trend for the GAD65+GAD67 protein levels (132.54±16.00; ns) in *Magel2*-deficient mice. At P30 (Figure 10d and 10e), we continued to observe a significant increase of *Gad65* (1.40±0.21; p < 0.01), *Gad67* (1.63±0.34; p < 0.01) transcripts, and the protein levels of GAD 65+GAD67 (142.31±19.15; p < 0.05). These results show an increase of expression of *Gad65* and *Gad67* at P5, which is more significant at P30 in *Magel2^-/-^* mice, suggesting a higher number of GABAergic neurons or an increase of GABA production and GABA activity in *Magel2*-deficient mice.

**Fig 10:**
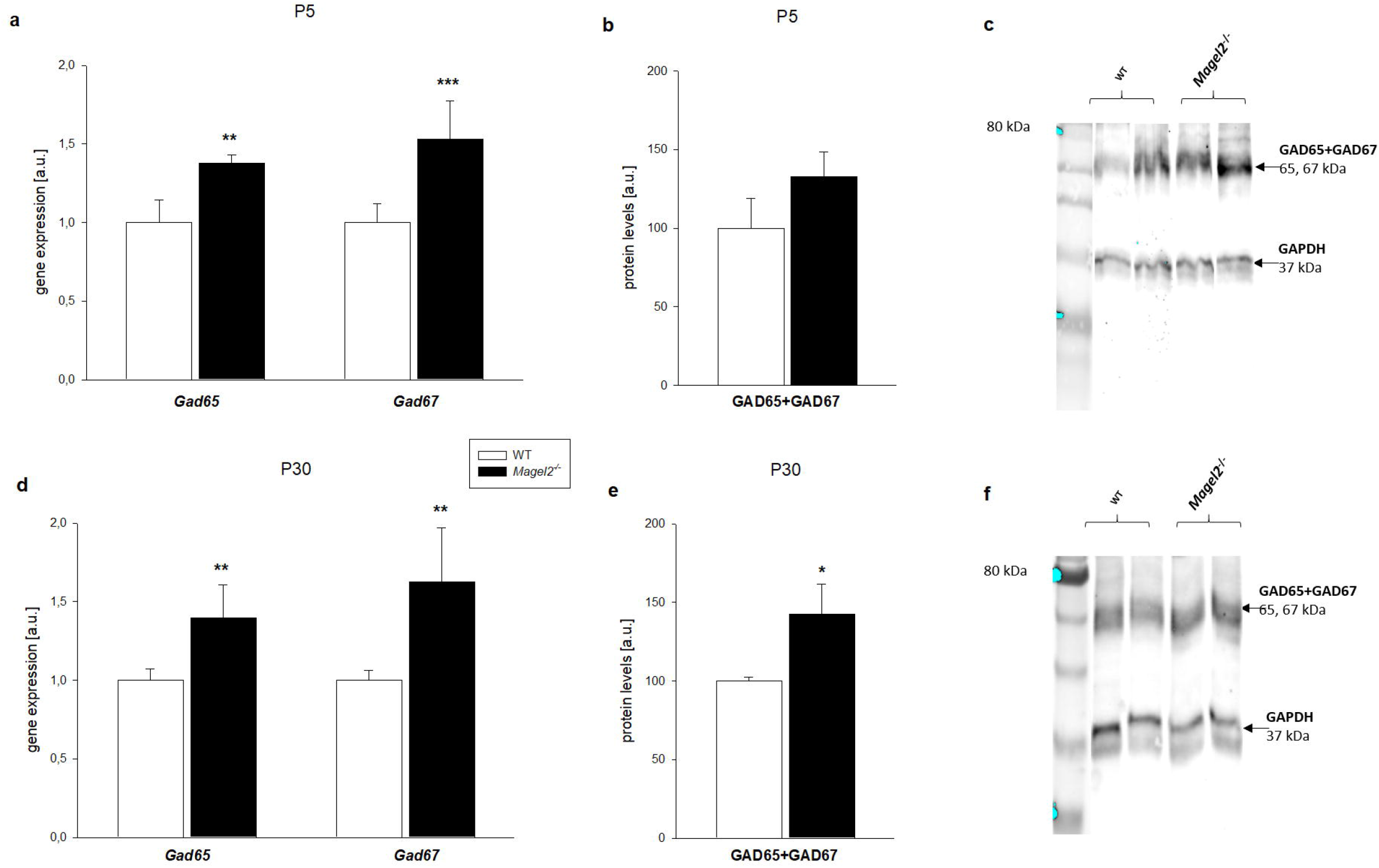
Changes in the hippocampal *Gad65* and *Gad67* transcript and GAD65+ GAD67 protein levels at P5 (a-c) and P30 (d-f) WT or *Magel2^-/-^* mice. The graphs show relative mRNA expression levels **(a, d)** normalized to *Gapdh* transcript and protein levels **(b, e)** normalized to GAPDH protein. qRT-PCR was calculated by the 2^-ΔΔCt^ Livak method (Livak & Schmittgen, 2001). Representative set of Western Blot Analysis results from GAD65+GAD67 at P5 **(c)** and P30 **(f)**. Means are represented on bars ± SEM (n = 4-6/group). At P5, the mRNA levels of *Gad65* (**p < 0.01, df=9, T=4.47) and *Gad67* (***p < 0.001, df=9, T=6.00) were significantly increased in *Magel2^-/-^* compared to WT mice. At P30, *Gad65* (**p < 0.01, df=10, T=4.43), *Gad67* (** p < 0.01, df=10, T=4.41) gene expression levels and protein level of GAD65+GAD67 (* p < 0.05, df=11, T=2.37) increased in *Magel2^--^* mice compared to the WT mice. *Glutamic acid decarboxylase - Gad*.

## Discussion

### Impaired neurite outgrowth in immature hippocampal neurons is accompanied by alteration in the retinoic acid signaling pathway in *Magel2*-deficient mice

Here, we showed impaired neurite outgrowth and dendritic arborization in primary cultures of immature hippocampal neurons isolated from *Magel2^-/-^* mice. Our data observed in hippocampal neurons are consistent with abnormal development of the hypothalamic neural projections at P8 *Magel2*-deficient mice found in the previous study by Maillard et al. (2016). Furthermore, the authors observed the role of *Magel2* in axonal growth by quantification of neurites in N2a cells transfected with *Magel2*. Moreover, a recent study has demonstrated impaired dendrite formation in induced pluripotent stem cell (iPSC)-derived neurons from Schaaf-Yang syndrome suffering patients (Crutcher et al., 2019). Interestingly, we found that ATRA failed to induce neurite extension in neurons isolated from *Magel2^-/-^* mice, although ATRA enhanced neurite outgrowth in the primary neurons of WT mice. ATRA, a naturally occurring retinoid metabolite of vitamin A, is routinely used as a differentiation factor, as well as a positive stimulus for the examination of neurite outgrowth (Clagett-Dame, McNeill & Muley, 2006; Kovalevich & Langford, 2013). Our data suggest that the retinoid signaling pathway is altered in *Magel2*-deficient animals. One possible explanation is offered by interference with the mTORC1 signaling pathway. It has been shown that mTORC1 is upregulated in the brains of *Magel2* null mice and accompanied by impairment of dendrite formation (Crutcher et al., 2019). Therefore, insufficient ATRA action on neurite outgrowth in *Magel2*-deficient mice could be associated with potential downstream alterations in vesicle trafficking, as well as the corresponding increased lipid droplet accumulation and changes in neuronal morphology observed in the study by Crutcher at al. (2019). Interestingly enough, it has been reported that ATRA can stimulate the expression of *Mage* family genes, including the *Magel2* in various cell lines *in vitro* (Gordeeva, Gordeev & Khaydukov, 2019), suggesting that *Magel2* is downstream of the ATRA pathway. In this study, we observed decreased expression of *Limk1* and *Tiam1*, which are enzymes known to be associated with rearranging of the cytoskeleton (see scheme in figure 11) and interact with the retinoid receptor function (Ishaq et al., 2011). Activation of LIMK1 is associated with retinoic acid-stimulated neurite outgrowth (Arastoo et al., 2016). LIMK1 regulates actin dynamics by phosphorylating cofilin, which contributes to processes of neurite elongation (Reichova et al., 2018). Since neurite elongation is essentially associated with functional intracellular transport processes, it is important to note that *Magel2* serves as a ubiquitin ligase enhancer and mediates the posttranslational modification of many substrates. One of the relevant issues regarding the impairments of neurite outgrowth is the mechanistic link of *Magel2* to the molecular machinery contributing to retrograde transport pathways (Hao et al., 2013).

### Hippocampal glutamatergic synaptic markers are decreased in *Magel2*-deficient mice

In primary hippocampal neurons, we found that *Magel2*-deficient mice have a lower amount of the presynaptic excitatory VGLUT2 transporter density. In addition, at day P5, the immunohistochemistry of PSD95/VGLUT2 colocalization showed a lower signal in the hippocampal areas CA1 and CA3 in *Magel2*-deficient mice. This lower density of the signal indicates that the number of glutamatergic neurons/synapses could be affected at this stage of development. Although glutamatergic synapses during the first postnatal week of development are not yet fully developed (Naskar et al., 2019), glutamatergic activity is necessary for their maturation. With regards to the demonstrated impairment of neurite outgrowth in the hippocampal cells in *Magel2*-deficient mice, it could be assumed that there is a delay in the maturation of hippocampal circuits. Under normal conditions, there are known synaptic connections between pyramidal neurons and interneurons within and between the areas of CA1 and CA2/CA (Booker & Vida, 2018). Indeed, in line with those observations, our other recent study has shown that the GABAergic activity was increased, and glutamatergic activity was reduced in the aCA3d pyramidal neurons when using hippocampal acute slices of Magel2-/- juvenile mice (Bertoni et al., 2020). It is assumed that the reduced glutamatergic activity most likely results from a postsynaptic alteration. The development of functional glutamatergic synapses could be delayed in conditions of *Magel2* deficiency. Interestingly, we also observed a higher expression of “GAD65+GAD67”, which suggests that *Magel2* deficiency at both developmental stages P5 and P30 is associated with enhanced GABAergic neurotransmission. In line with that, we previously observed an increase of GABAergic activity in *Magel2*-deficient hippocampal neurons (Bertoni et al., 2020). Indeed, it appears that *Magel2* deficiency is associated with GABAergic and glutamatergic alterations. Our results indicate more profound alterations at P5 compared to P30 *Magel2*-deficient mice, thus suggesting a delay in neuronal and circuitry maturation.

### Deficits observed in *Magel2*-deficient mice are reversed by OT treatment

We found that OT increased the neurite length in the WT and *Magel2^-/-^* primary hippocampal neurons. This is in line with our previous results on neuronal cell lines treated with OT (Lestanova et al., 2016; Zatkova et al., 2018). Although it has been demonstrated that *Magel2* - deficient neonates suffer from a significant reduction in OT (Schaller et al., 2010), OXTRs could be fully functional, which explains the stimulatory effect of OT on *Magel2^-/-^* neurons. Globally, in the hippocampus at P5, we observed an increase of expression of several transcripts involved in neurite outgrowth following an administration of OT (suggesting a stimulatory role of OT in these processes). In addition, we showed that OT administration reduces the quantity of PSD95/VGLUT2 colocalization in the WT mice with no additional effect in *Magel2*-deficient mice. Thus, we suggest that when neurons are still immature, OT administration reduces the quantity of glutamatergic synapses in the hippocampal regions of WT mice; however, not in *Magel2*-deficient mice, which had previously presented decreased glutamatergic activity. In line with our results, a recent study showed that OT treatment controls the development of hippocampal glutamatergic neurons (Ripamonti et al., 2017). In the context of pathophysiological conditions present in *Magel2*-deficient mice, an alteration in OT system might result in abnormal and delayed development of hippocampal circuits. In correspondence to this, another study involving adult *Magel2*-deficient mice has found reduced excitatory and increased inhibitory currents in OT neurons (Ates et al., 2019). We need to be careful, however, when making any conclusions, since the role of OT could be specifically-related to the developing immature hippocampal neurons and could also depend on the density of OXTRs in specific hippocampal subregions (Raam et al., 2017; Young & Song, 2020).

## Conclusion

This is the first time that an alteration in the neurite outgrowth of immature hippocampal neurons in *Magel2^-/-^* mice has been demonstrated to be accompanied by a decrease in the expression of factors associated with neurite growth. We found that OT compensates for the dendritic arborization defects observed in primary *Magel2^-/-^* hippocampal neurons and stimulates the expression of factors involved in neurite outgrowth overall (see scheme in figure 11). A deficit of glutamatergic synapses has also been shown in *Magel2*-deficient mice that are not restored after OT administration. Unexpectedly, our results suggest that OT administration in WT mice induces a reduction in the number of glutamatergic synapses, supporting the idea that OT controls the formation of glutamatergic synapses. These results reveal the strong impact of early childhood OT treatment on the development of the nervous system and the interest in certain neuropsychiatric conditions.

## Acknowledgement

We would like to thank Dr. Karel Frimmel, Centre of Experimental Medicine SAS for his assistance with confocal microscopy, Dr. Igor Medina, INMED, Marseille for critical reading, as well as Michael Sabo for proofreading of the manuscript.

## Data availability statement

The data that support the findings of this study are available from the corresponding author upon reasonable request.

## Author Contributions

JB, FM, ZB, and AR designed the research. AR, FS and SB performed the research. ZB, AR and JB analyzed the data. AR, FM and JB wrote the paper.

## Abbreviations

ASD: autism spectrum disorder
ATRA: all-trans retinoic acid
DIV: day *in vitro*
GABA: γ-aminobutyric acid
GAD: glutamic acid decarboxylase
GAP43: growth associated protein 43
GAPDH: glyceraldehyde 3-phosphate dehydrogenase
LIMK1: LIM domain kinase 1
MAGEL: MAGE Family Member L2
MAP2: microtubule associated protein 2
mTORC1: mechanistic (or mammalian) target of rapamycin complex 1
NLGN: neuroligin
NRXN: neurexin
OT: oxytocin
OXTR: oxytocin receptors
PAK1: p21 activated kinase 1
PSD95: postsynaptic density protein 95
PWS: Prader–Willi Syndrome
Rac1: Ras-related C3 botulinum toxin substrate 1
RhoA: Ras homolog family member A
RhoB: Ras homolog family member B
SHANK: SH3 and multiple ankyrin repeat domains
SYS: Schaaf-Yang Syndrome
TIAM1: T-cell lymphoma invasion and metastasis 1
TRIM27: tripartite motif containing 27
USP7: ubiquitin specific peptidase 7
VGLUT2: vesicular glutamate transporter 2
WASH: Wiskott–Aldrich syndrome protein and SCAR homologue
WT: wild type

---

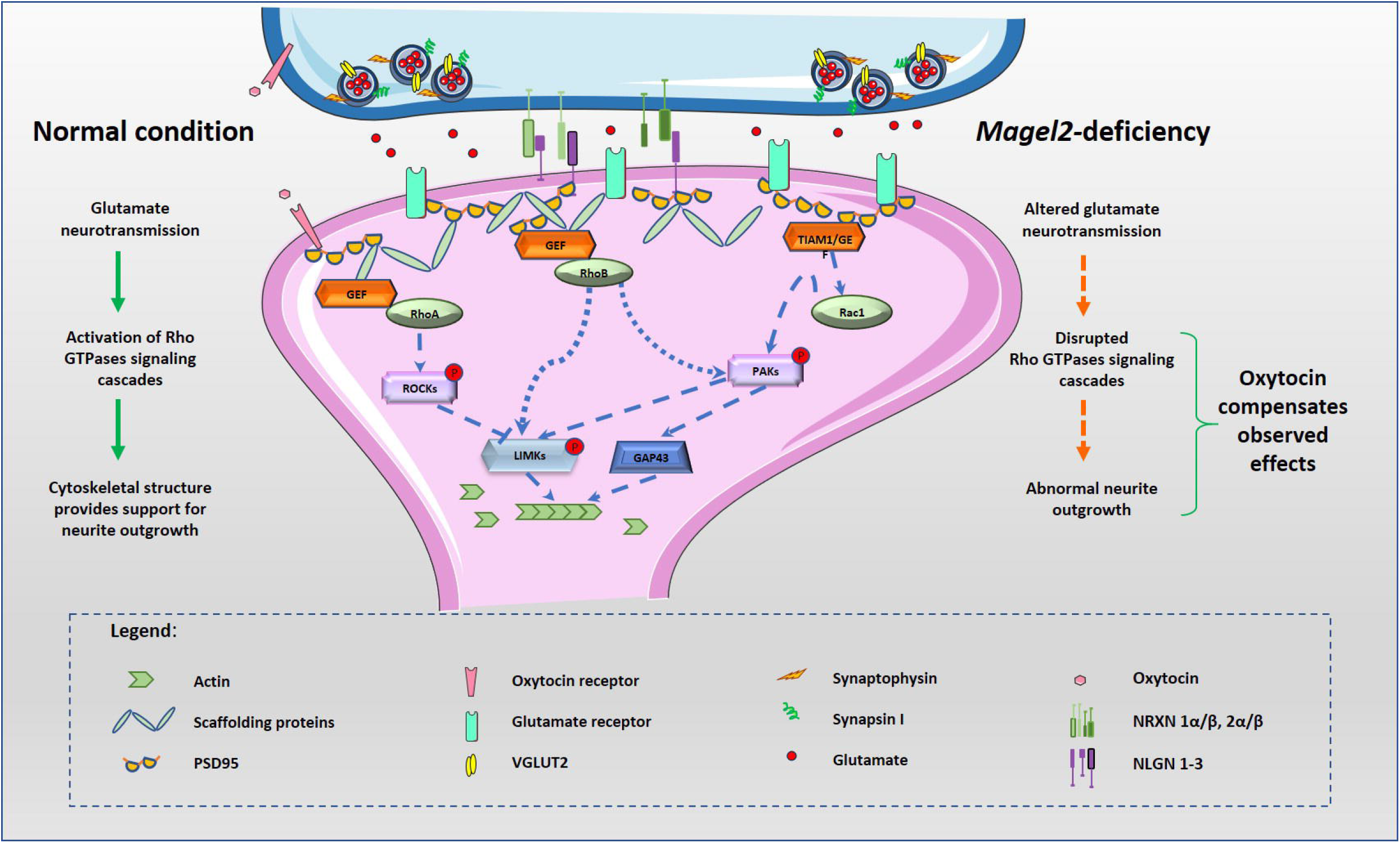

